# Repurposing of the multiciliation gene regulatory network in fate specification of Cajal-Retzius neurons

**DOI:** 10.1101/2022.11.18.517020

**Authors:** Matthieu X Moreau, Yoann Saillour, Vicente Elorriaga, Benoît Bouloudi, Elodie Delberghe, Tanya Deutsch Guerrero, Amaia Ochandorena-Saa, Laura Maeso-Alonso, Margarita M Marques, Maria C Marin, Nathalie Spassky, Alessandra Pierani, Frédéric Causeret

## Abstract

Cajal-Retzius (CR) neurons are key players of cortical development that display a very unique transcriptomic identity. However, little is known about the mechanisms involved in their fate specification. Here we use scRNAseq to reconstruct the differentiation trajectory of hem-derived CR cells (CRs) and unravel the transient expression of a complete gene module previously known to control the cellular process of multiciliogenesis. However, we find that CRs do not undergo centriole amplification or multiciliation. We show that upon genetic disruption of *Gmnc*, the master regulator of the multiciliation cascade, CRs are initially produced but fail to reach their normal identity and lean towards an aberrant fate resulting in their massive apoptosis. We further dissect the contribution of multiciliation effector genes and identify *Trp73* as a key determinant. Finally, we use *in utero* electroporation to demonstrate that the intrinsic competence of hem progenitors as well as the heterochronic expression of *Gmnc* prevent centriole amplification in the CR lineage. Our work exemplifies how the co-option of a complete gene module, repurposed to control a completely distinct process, may contribute to the emergence of novel cell identities.

## INTRODUCTION

Cajal-Retzius (CR) cells are transient glutamatergic neurons of the developing forebrain that reside in the marginal zone until their elimination by apoptosis during the first postnatal weeks in rodents (Causeret et al., 2021). Since their discovery more than a century ago by S. Ramon y Cajal and G. Retzius, they have been shown to play a fundamental role in the control of cortical lamination, arealization and circuit formation via the secretion of signalling factors, including Reelin. CR cells (CRs) presence is currently reported in amniotes only, suggesting they represent a relatively recent evolutionary innovation, and their abundance in the mammalian dorsal pallium led to propose they could have contributed to brain complexification during evolution (Goffinet, 2017).

In mouse, CRs are generated between embryonic day (E) 10.5 and E13.5 (Hevner et al., 2003; Takiguchi-Hayashi et al., 2004) in discrete pallial progenitors domains located at the border of the cortical primordium, from which they migrate tangentially in the marginal zone to cover the entire cortical surface. Genetic tracing experiments allowed the identification of four distinct ontogenic origins for CRs, three of which course along the pallial midline: the septum rostrally (Bielle et al., 2005), the cortical hem that produces the majority of neocortical CR cells (Takiguchi-Hayashi et al., 2004; Yoshida et al., 2006), and the thalamic eminence (ThE) caudally (Ruiz-Reig et al., 2017; Tissir et al., 2009). An additional described origin corresponds to the pallial-subpallial boundary (PSB), also referred to as ventral pallium (Bielle et al., 2005). We have previously shown that hem-, septum- and ThE-derived CR cells (subsequently referred to as medial CRs) share the expression of a highly specific gene module comprising *Trp73, Lhx1, Cdkn1a* (p21), *Ebf3* and *Cacna2d2*, among other genes, distinguishing them from PSB-derived CRs that lack such signature (Moreau et al., 2021). Yet, all subtypes share common features: they are the only *Foxg1*-negative neurons in the developing cerebral cortex and additionally express high levels of *Reln* and *Nhlh2* (Moreau et al., 2021). In *Foxg1*-deficient mice, cortical neurons abnormally express Reln, suggesting that CRs represent the default fate unless repressed by Foxg1 (Hanashima et al., 2004). However, only *Trp73*^-/-^ CRs are produced upon conditional inactivation of *Foxg1* at E13 (Hanashima et al., 2007), pointing to additional mechanisms involved in the specification of medial CRs. To date, such mechanisms remain completely elusive.

Interestingly, medial CR markers *Trp73* and *Cdkn1a* were recently involved in multiciliation (Marshall et al., 2016; Ortiz Álvarez et al., 2021), an evolutionary ancient cellular process consisting in the amplification of centrioles followed by growth of multiple motile cilia. The central nervous system of all vertebrates is lined by multiciliated ependymal cells, and multiciliated cells of the choroid plexus (ChP) secrete the cerebrospinal fluid (Lun et al., 2015; Spassky and Meunier, 2017). The gene regulatory network controlling multiciliation is also highly conserved between organs and species. The chromatin regulator GemC1, encoded by *Gmnc*, is both necessary and sufficient for multiciliation, and was shown to control the expression of most major components of the multiciliation cascade, including *Trp73* and *Cdkn1a* (Kyrousi et al., 2015; Lalioti et al., 2019; Ortiz Álvarez et al., 2021; Terré et al., 2016).

In this study we used single-cell RNA sequencing (scRNAseq) to investigate the mechanisms involved in the specification of medial CRs, focussing on the most abundant source located at the hem. We found that all components of the multiciliation gene regulatory network are expressed transiently at early steps of CR differentiation. Yet, CRs do not amplify their centrioles or grow multiple cilia. We then disrupted the multiciliation regulatory network to demonstrate that *Gmnc* is required for CR specification. In its absence, CRs are produced, acquire some of their cardinal features, but fail to maintain their identity and undergo cell death. *Trp73* mutants completely phenocopy *Gmnc*-deficient animals, unlike *Ccdc67/Deup1, Ccno* or *Ccdc40* mutants, pointing to a differential requirement of multiciliation genes in CRs specification. Using *in utero* electroporation, we demonstrate that competence of the hem territory as well as heterochrony in *Gmnc* expression restrict its ability to induce multiciliation. Our work exemplifies how an evolutionary ancient gene network can be coopted and repurposed to perform a function completely distinct from its original one.

## RESULTS

### Transcriptomic characterization of the early medial telencephalon

In mouse, the cortical hem is the main source of neocortical CRs, with a contribution estimated between 60% and 90% (Yoshida et al., 2006). In order to better understand the process of CR specification, we performed scRNAseq on dissected tissue from the dorsal midline, encompassing the hem **(Figure 1A)**. To ensure sufficient sampling of CR progenitors, precursors and neurons throughout the specification and early differentiation processes, we pooled tissue obtained from four E11.5 and four E12.5 embryos. E11.5 samples were collected from *PGK^Cre^;Rosa26^YFP^* embryos and E12.5 cells were obtained from *Wnt3a^Cre^;Rosa26^tdTomato^* animals, to enable the deconvolution of each stage on the basis of *YFP* sequencing reads, and the identification of E12.5 hem and hem-derived cells thanks to *Tomato* expression (**Figure S1A**).

**Figure 1.**
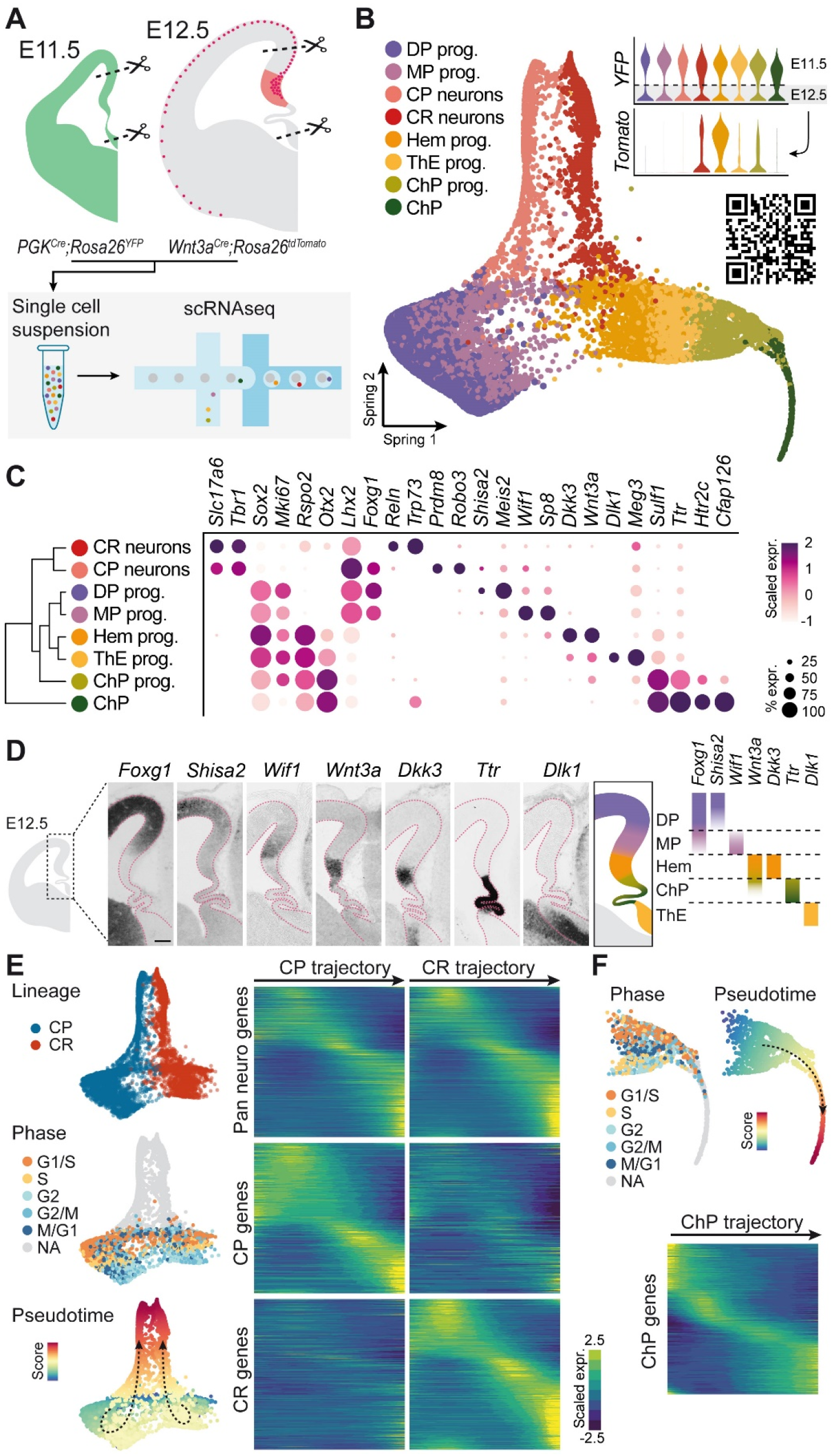
Transcriptional landscape around the early mouse hem. (A) The hem region from E11.5 *PGK^Cre^;Rosa26^YFP^* and E12.5 *Wnt3a^Cre^;Rosa26^tdTomato^* mouse embryos was dissected and subjected to scRNAseq. (B) 2D manifold representation of the dataset obtained after SPRING dimensionality reduction. Violin plots depict the contribution of *YFP*^+^ and *Tomato*^+^ cells to each cluster. The QR code points to the Shiny App allowing to browse the data. (C) Bubble plot showing differential gene expression between clusters. The dendrogram indicates the hierarchy between clusters. (D) ISH on coronal sections of the E12.5 forebrain for selected genes indicating the position of each domain. Scale bar: 100μm. The diagram on the right uses the same color code as in (B). (E) Reconstruction of CRs and CP differentiation trajectories (left). Heatmaps (right) represent gene expression along pseudotime for each lineage, using gene sets common or specific to each trajectory. (F) Reconstruction of ChP differentiation trajectory and identification of gene differentially expressed along pseudotime. DP: dorsal pallium, MP: medial pallium, CP: cortical plate, CR: Cajal-Retzius, ThE: thalamic eminence, ChP: choroid plexus.

After quality control and filtering, we obtained 15,160 cells with a median 3,750 genes detected per cell which were subject to iterative clustering steps and annotation based on known marker genes expression. Projection of these cells on a low-dimensional SPRING embedding (**Figure 1B**) revealed 5 clusters containing cycling progenitors, collectively identified by the expression of *Sox2* and *Ki67* (**Figures 1B, C and S1B**). Dorsal pallium (DP) and medial pallium (MP) progenitors, both *Foxg1^+^/Lhx2^high^*, were distinguished by the expression of *Shisa2/Meis2* and *Wif1/Sp8*, respectively (**Figures 1C and S1B**). They segregated away from the hem, ThE and ChP progenitors, all *Rspo2*^+^/*Otx2*^+^ and best identified by the expression of *Dkk3*/*Wnt3a, Dlk1/Meg3* and *Sulf1/Ttr*, respectively. Tissue mapping of these progenitor domains by *in situ* hybridization (ISH) highlighted their spatial segregation and confirmed the sharp distinction between the abutting medial pallium and hem (**Figures 1D and S1B, C**). Three postmitotic differentiation trajectories emerged from those progenitors: two of them contained pallial glutamatergic neurons (*Tbr1*^+^/*Slc17a6*^+^), and corresponded to cortical plate (CP) neurons (*Prdm8^+^*/*Robo3^+^*) and CR cells (*Trp73*^+^/*Reln^high^*), whereas the remaining one was non-neuronal and were identified as ChP (*Htr2c*^+^/*Cfap126*^+^). *YFP*^+^ cells (collected at E11.5) were found in all clusters with a contribution of ~65 %, showing no strong cell composition bias between stages (**Figure S1A**), consistent with the idea that the cellular processes ongoing at E11.5 and E12.5 do not significantly differ. *Tomato*^+^ cells (belonging to the Wnt3a lineage) were most abundant in the hem, CR and ChP progenitors clusters (**Figure 1B**), consistent with previous reports (Yoshida et al., 2006). Of note, the few *Tomato*^+^ cells found in the ThE cluster rather reflect imperfect clustering than the real presence of Wnt3a-derived cells in this tissue (not illustrated).

We then attempted to reconstruct the transcriptional trajectory followed by CP neurons, CR cells and ChP throughout the specification and early differentiation steps. The parallel arrangement of the two neuronal trajectories suggest they share the sequential expression of pan neuronal transcriptional modules and differ by sets of lineage-specific genes. CR and CP neurons, together with their progenitors of origin, were aligned along a pseudotime axis to extract such common or lineage-specific genes and characterize their expression dynamics during fate commitment and early differentiation (**Figure 1E**). A similar strategy was used to characterize the transcriptional dynamics during ChP differentiation (**Figure 1F**). Importantly, the high cellular sampling in each lineage allowed to capture the successive transient cell states at relatively high resolution. The approach was validated by the observation that pan-neuronal genes such as *Sox2, Neurog2* and *Tbr1* were sequentially expressed with similar dynamics in the two neuronal lineages, whereas *Wnt3a, Trp73* and *Lhx1* or *Tgfb2, Rprm* and *Prdm8* were specifically and sequentially expressed along CR and CP trajectories, respectively (**Figure S1D**).

We therefore generated a high-resolution transcriptomic atlas of the hem and surrounding regions of the early developing telencephalon, including plain differentiation trajectories for CR, CP and ChP lineages. In addition, we provide a web-based userfriendly Shiny App interface to explore the data (available at https://apps.institutimagine.org/mouse_hem/).

### CR cells transiently express multiciliation genes but are not multiciliated

Since mature CRs harbour a very peculiar transcriptomic signature compared to other cortical excitatory neurons (Causeret et al., 2021; Moreau et al., 2021; Tasic et al., 2018), we speculated they should express lineage-specific fate determinants at the onset of their differentiation. We therefore focused on the set of genes upregulated upon commitment to neuronal differentiation in the CR lineage but not in CP neurons. We identified 92 genes specifically expressed at the exit of the apical progenitor state in the CR lineage (**Figure 2A**). In order to identify pathways possibly involved in CR fate specification, we subjected the gene list to a GO-term enrichment analysis which surprisingly revealed a significant over-representation of genes associated with the differentiation of multiciliated cells (**Figure 2B**). We therefore computed a multiciliation signature, consisting in 28 genes currently known to regulate this process (Lewis and Stracker, 2021), to find out that their expression during CR differentiation is transiently upregulated at levels matching that of ChP (**Figure 2C**). By contrast, CP neurons did not display any increase in the expression of the multiciliation gene module throughout differentiation. We then wondered what could be the extent of transcriptomic similarity between differentiating CRs and ChP. We found that among the 612 genes specific of the CR trajectory, more than half (319) were shared with ChP **(Figure 2D**). This indicates a major overlap between the differentiation programs at play in both cell types, extending well beyond the few dozens of known multiciliation genes. Nevertheless, CRs and ChP each displayed strong expression in pallial progenitors is sufficient to induce centriole amplification, leading to the growth of multiple cilia and acquisition of an ependymal fate (Kyrousi et al., 2015), prompted us to investigate whether CRs could undergo multiciliation during their specific signatures, consistent with their expected distinct specialisation.

**Figure 2.**
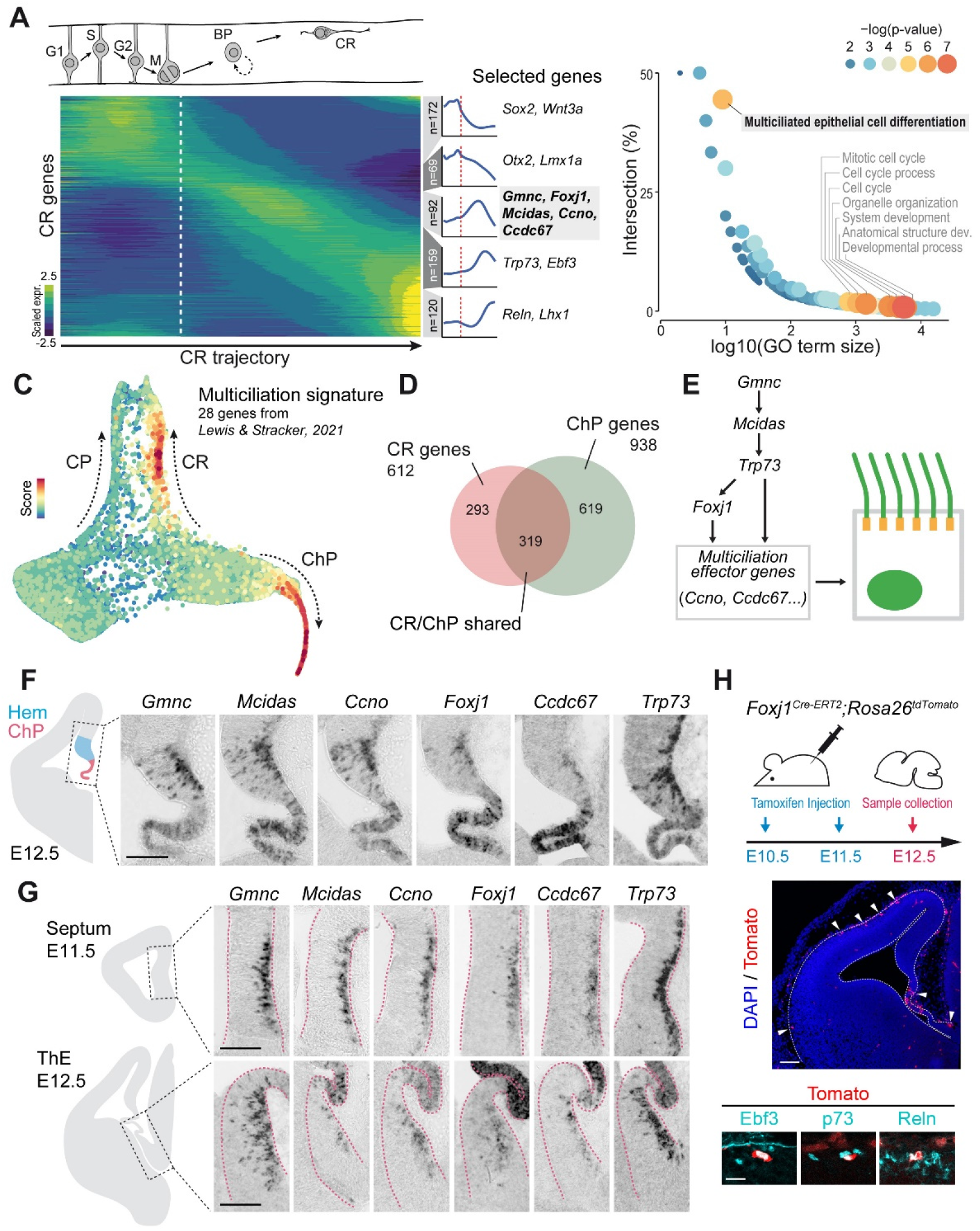
Expression of multiciliation genes in differentiating CR cells. (A) Heatmap depicting gene expression during CRs differentiation. The average trends of the successive gene waves are shown on the right. Selected genes characteristic of each wave are indicated. Dashed lines mark the exit of apical progenitor state. (B) GO-term analysis revealing the statistical enrichment of multiciliation-related genes during CRs differentiation. (C) SPRING representation showing that the average expression of 28 known multiciliation genes is transient during CRs differentiation and reaches levels similar to ChP. (D) Venn diagram highlighting the important overlap between CRs- and ChP-enriched genes. (E) Simplified view of the gene regulatory network involved in centriolar amplification and multiciliation. (F-G) ISH on coronal sections of the E11.5 or E12.5 forebrain validating the expression of multiciliation genes in the hem (F), septum and ThE (G). (H) Genetic tracing followed by immunostaining indicate that the progeny of *Foxj1-* expressing cells (arrows) is found in the ChP, ThE and neocortical marginal zone. The latter express CRs markers Ebf3, p73 and Reln. Scale bars: 100μm in F, G and H (top image), 20μm in H (bottom panel).

At the individual gene level, we found that the main components of the multiciliation network (**Figure 2E**) exhibit distinct expression dynamics, with the master regulator *Gmnc* starting at the very onset of the CR trajectory and preceding all other genes, as exemplified with *Mcidas, Ccno, Foxj1, Ccdc67/Deup1* and *Trp73* (**Figure S2A,B**). ISH for these genes revealed their shared expression at all sources of medial CRs, as signal was observed in the hem (**Figure 2F**) as well as the septum and ThE (**Figure 2G**), contrasting with the PSB in the lateral aspect of the cortex where we never detected expression (not illustrated). All multiciliation genes, to the exception of *Trp73*, were expressed transiently in differentiating CRs, showing a strong downregulation when they engage tangential migration in the neocortex.

To formally confirm those findings, we crossed the *Foxj1^Cre-ERT2^* mouse line with the *Rosa26^tdTomato^* reporter strain and perform tamoxifen injections at E10.5 and E11.5 to permanently label the early-born progeny of *Foxj1* -expressing cells. Embryos collected at E12.5 showed numerous Tomato^+^ cells occupying the characteristic superficial position of CRs in the developing cortex and immunoreactive for CR markers Ebf3, p73 and Reln (**Figure 2H**). This experiment confirms that CRs are generated from *Foxj1*-expressing progenitors, consistent with recent data using a distinct *Foxj1^Cre^* strain (Parichha et al., 2022).

We therefore conclude that CR cells originating from medial sources (hem, septum and ThE) transiently express an apparently complete gene module known to regulate multiciliation at the onset of their differentiation.

We then decided to investigate whether the expression of multiciliation genes in differentiating CR cells was evolutionarily conserved. Indeed, CR cells have been postulated as important players during brain evolution (Goffinet, 2017), and the presence and diversity of CR subtypes in the avian pallium remains a matter of debate (Causeret et al., 2021). We detected strong expression of *Gmnc* in the septum and hem regions of E5 chick embryos (Figure **S2C**), in direct apposition to cells expressing *Trp73, Ebf3* and *Reln* located in the marginal zone. We conclude that the transient expression of multiciliation genes during the differentiation of CR cells originating from medial sources is most likely evolutionarily conserved among amniotes.

So far, CRs have not been reported to be a multiciliated cell type. However, previous work showing that *Gmnc* differentiation. We thus performed immunostaining for the centriole marker FOP on E12.5 embryos. Newly generated CR s, identified by their position in the hem and expression of p73 displayed a single FOP^+^ foci, sometimes sufficiently resolved to distinguish a pair of centrioles (**Figure 3A, B**). In addition, flat-mount neocortical preparation of *Wnt3a^Cre^;Rosa26^tdTomato^* embryos indicated that genetically-identified CR cells that had migrated away from the hem in the dorsal pallium also bear a single pair of centrioles, contrasting with ChP from the same sample that showed robust centriolar amplification (**Figure 3C, D**). Finally, Arl13b-positive cilia above p21^high^ CR nuclei in the hem appeared well defined, short and straight (**Figure 3E**), whereas their counterparts in p21^low^ ChP were found long, tortuous and intermingled (**Figure 3F**). Thus, while CRs express transiently all the major known multiciliation genes, they do not undergo centriole amplification or multiciliation upon differentiation.

**Figure 3.**
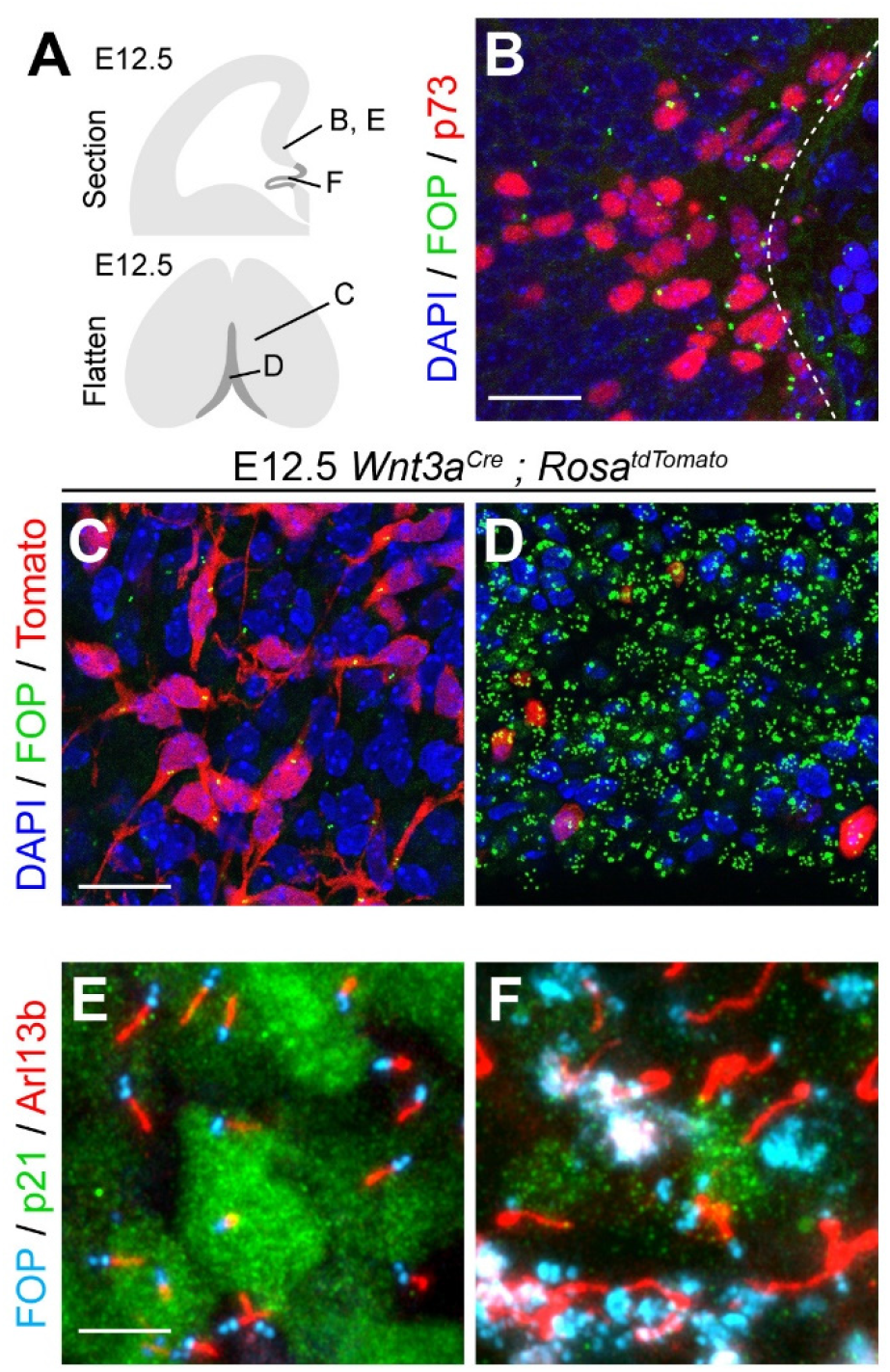
Centrioles and cilia in CR cells and ChP. (A) Drawing indicating the stage, position and preparations illustrated in (B-F). (B) Immunofluorescence showing a single centriole (FOP^+^, green) in nascent CRs (p73^+^, red) of the hem. Nuclei are stained with DAPI (blue). (C-D) Immunostaining of the neocortical pial surface (C) and ChP ventricular surface (D) showing centrioles (FOP^+^, green) and cells derived from *Wnt3a*-expressing progenitors of the hem (Tomato^+^, red), corresponding to CRs cells (C) as well as rare ChP cells (D). Nuclei are stained with DAPI (blue). (E-F) Immunostaining of the hem (E) and ChP (F) using antibodies against FOP (cyan) and Arl13b (red) to visualize centrioles and cilia, respectively. Immunoreactivity for p21 is strong in CRs (E) and weaker in ChP (F). Scale bars: 20μm in B and C, 10μm in E.

### The multiciliation master gene *Gmnc* is required for medial CR fate specification

The absence of centriolar amplification in CRs raises the question of the functional implication of the multiciliation gene module during CR differentiation. We therefore investigated the consequences of a complete disruption of the multiciliation regulatory network, using mouse deficient for *Gmnc*, the gene described as the most upstream regulator of the cascade (Lewis and Stracker, 2021) (**Figure 2E**).

We first performed scRNAseq on E12.5 hem explants collected from *Gmnc^-/-^* embryos. After quality control and filtering, we obtained 10,036 cells with a median 3,123 genes detected per cell. SPRING dimensionality reduction revealed a topology reminiscent of the wild-type dataset, with *Sox2^+^/Foxg1^+^* MP/DP progenitors segregated away from *Wnt3a^+^* hem, *Dlk1^+^* ThE and *Ttr*^+^ choroid domains (**Figure S3A**). Neuronal trajectories were also readily identified, with the presence of *Tbr1^+^* neurons emerging from both MP/DP and Hem/ThE progenitors, suggesting that neuronal production is not affected in the absence of *Gmnc*, although they appear less obviously resolved than in the wild-type condition. Consistent with previous data (Terré et al., 2016), we confirmed that *Gmnc* target genes, including *Mcidas, Ccno, Ccdc67/Deup1* and *Trp73* were absent from the mutant dataset, whereas transcripts from the mutant *Gmnc* locus were still detectable as they retained the 3’UTR (**Figure S3B**). In order to better appreciate changes upon *Gmnc* loss, we performed cell type label transfer to predict progenitors and ChP identities in the mutant dataset using the wild-type as a reference, as well as dataset integration and projection of mutant cells onto the wild-type reference (**Figure 4A**). Whereas pallial, hem, ThE and ChP progenitors were found and correctly represented in *Gmnc* mutants, not a single cell was annotated as differentiated ChP, consistent with previous reports indicating defective ChP multiciliation in *Gmnc* mutant animals (Terré et al., 2016). Yet, ChP were correctly specified as indicated by the normal expression of *Ttr* or *Htr2c* (**Figure S3C**). Furthermore, the wild-type CR differentiation trajectory remained unmatched by mutant cells that instead occupied a position close to CP neurons in the 2D embedding (**Figure 4A**). To circumvent such an apparent coalescence of CR and CP neuronal trajectories in the mutant dataset, we isolated mature neurons, subjected them to unsupervised hierarchical clustering and reconstructed their lineage of origin using FateID (**Figure S3D**). The smallest of the three identified lineages contained the only *Lhx1*^+^ cells remaining in the mutant dataset, and was also characterized by the expression of ThE genes *Gm27199, Grm1, Chl1* and *Unc5c* (**Figure S3D-E**). Since *Lhx1* expression was found restricted to the ThE and ventral migratory stream in *Gmnc^-/-^* embryos (**Figure S3F**), we concluded these cells most likely corresponds to ThE-derived CRs. Because of their low sampling, we excluded this small subset from subsequent analyses. Among the two remaining neuronal lineages, CP neurons were readily distinguished as *Foxg1^+^/Prdm8^+^/Robo3^+^* (**Figure 4B**). The remaining population displayed a transcriptomic signature reminiscent of CR cells, characterized by the absence of *Foxg1* and high expression of *Reln*, *Lhx5, Nr2f2, Rspo3*, *Zic5 or Zfp503* (**Figure 4B** and data not shown), and were therefore annotated as such. These mutant CRs clearly diverged from their wild-type counterparts and appeared to progressively lean towards a CP fate, although not completely acquiring such identity as indicated by measurement of their cosine distance along pseudotime (**Figure 4C**).

**Figure 4.**
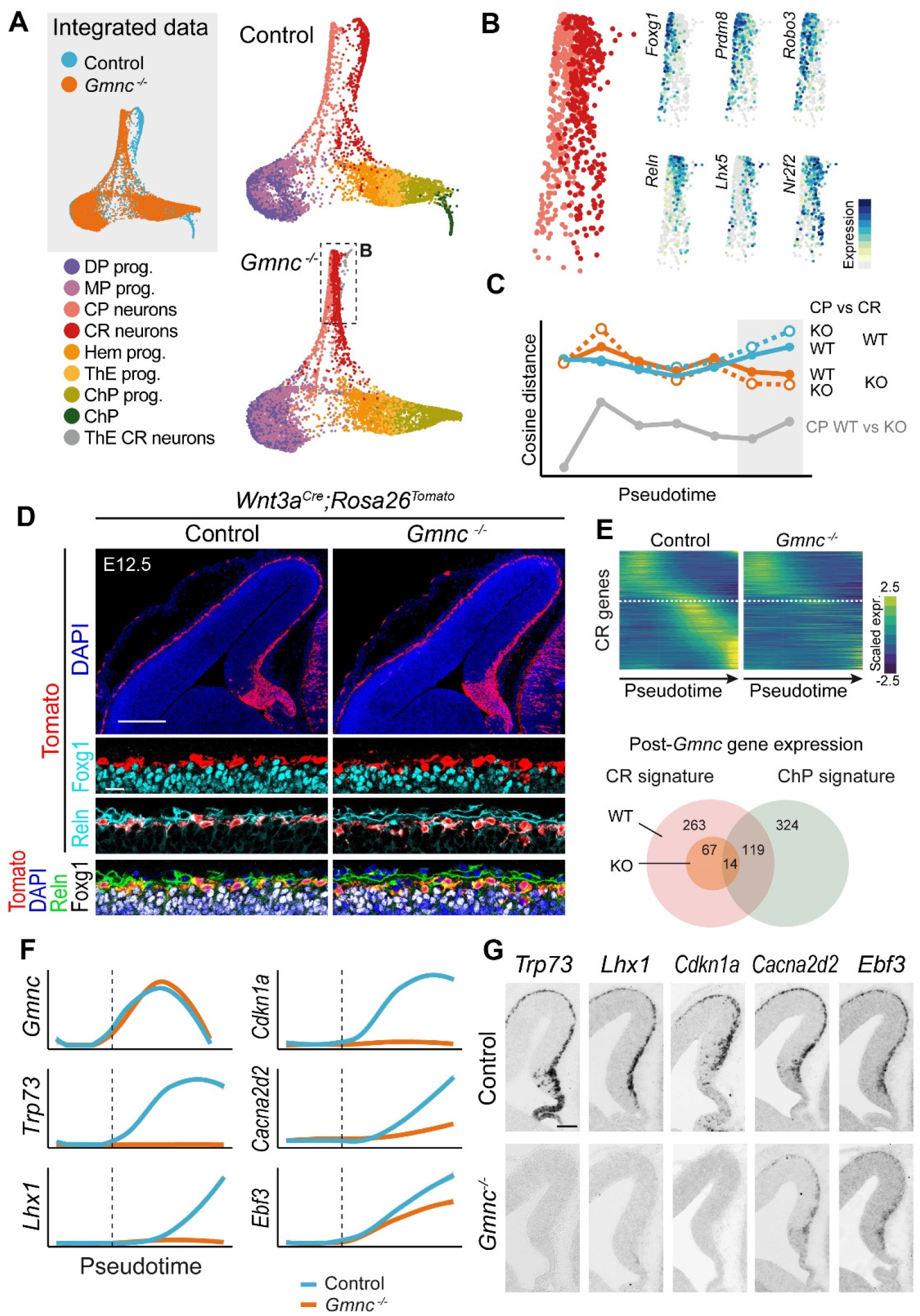
CR fate specification in *Gmnc* mutants. (A) SPRING embedding of integrated scRNAseq data from control embryos and *Gmnc* mutants. Cell type annotation of the mutant dataset was achieved by label transfer, using the control dataset as a reference. (B) High magnification of (A) indicating that mutant CRs can be distinguished from CP despite their apparent coalescence, as exemplified with the expression of selected genes. (C) Cosine distance between CRs and CP along pseudotime, the area in grey corresponds to the most differentiated states. Colors indicate the genotype of CR, whereas line types refer to the genotype of CP. CRs from *Gmnc* mutants (orange lines) differ less from CP than their WT relatives (blue lines), regardless of the genotype of CP. (D) Immunostaining on coronal sections of the pallium of E12.5 *Wnt3a^Cre^;Rosa26^tdTomato^* control or *Gmnc* mutant embryos. For both genotypes, hem-derived (Tomato^+^) cells are found in the neocortical marginal zone and are Foxg1^-^ and Reln^+^. (E) Heatmap (top) showing the strong disruption of CRs-specific gene expression during CRs differentiation in *Gmnc* mutants. The dashed line corresponds to *Gmnc*. Venn diagram (bottom) representing the decreased transcriptional overlap between CRs and ChP following *Gmnc* loss of function. (F) Temporal expression dynamics of selected genes during CRs differentiation in control (blue) and *Gmnc* mutant (orange) conditions. Mutant *Gmnc* transcripts lack exons 3 and 4 but retain the 3’UTR and are therefore detected by scRNAseq. The dashed lines corresponds to the exit of apical progenitor state. (G) ISH on coronal sections of E12.5 embryos confirming the complete loss, or severe reduction, of expression of the CRs marker genes shown in (F) in *Gmnc* mutants. Scale bars: 200μm in (D), 100μm in (G), 20μm in high magnifications in (D).

We then performed genetic tracing using the *Wnt3a^Cre^* driver in control and *Gmnc* mutant backgrounds at E12.5 to find out that, regardless of the presence or absence of *Gmnc*, hem-derived cells migrate tangentially to cover the entire neocortex, and remain Foxg1^-^ and Reln^+^, all canonical features of CR fate (**Figure 4D**). In addition, we found the neurogenic progression, illustrated by the sequential expression of *Sox2, Neurog2*, and *Tbr1*, similar between control and mutant CR lineages (**Figure S4A**). Tbr2 immunostaining of the hem also revealed no difference between genotypes (**Figure S4B**), further supporting the conclusion that *Gmnc* is dispensable for CR cell production. However, pseudotime analyses indicated that the transcriptional dynamics associated with CR identity was dramatically affected in mutants, as only a small fraction of the genes normally upregulated after the onset of *Gmnc* expression were still differentially expressed in the mutant CR lineage (81 out of 382, **Figure 4E**). The substantial overlap between CR and ChP-specific genes observed in wild-type animals was no longer present in *Gmnc* mutants (**Figure 4E**), indicating that most of the similarity between the two lineages corresponds to *Gmnc*-dependent genes. These results indicate that the multiciliation gene regulatory network is a strong contributor not only to ChP, but also CR identity. Consistently, classical marker genes of medial CRs, such as *Trp73, Lhx1* or *Cdkn1a*, were not detected in the mutant trajectory, whereas others like *Cacna2d2* or *Ebf3* displayed decreased expression (**Figure 4F**). ISH experiments confirmed these findings, showing a complete loss of *Trp73, Lhx1*, and *Cdkn1a* expression, and a strong reduction of *Cacna2d2* and *Ebf3* in the neocortex of mutants (**Figure 4G**). Conversely, we failed to identify genes convincingly upregulated in mutant CRs, suggesting that their apparent proximity with CP neurons is not due to fate convergence, but rather to a loss of the aforementioned medial signature. Taken together, these observations demonstrate a normal production but improper fate specification of CRs upon disruption of the multiciliation gene network.

### Establishment of a complete identity is required for CRs survival

To investigate the outcome of mis-specification on CR fate, we first performed ISH experiments at E13.5 in *Gmnc* mutants and control littermates. We observed a striking absence of expression of all tested marker genes of medial CR identity, not only those like *Trp73* that were already absent at E12.5, but also *Ebf3* or *Cacna2d2* that were still detected in mutants one day earlier (**Figures 5A-C and S5,** compare with **Figure 4G**). ISH for genes expressed by all CR subtypes (including those originating from the PSB) such as *Reln* or *Nhlh2* revealed an almost complete loss of signal in the dorsal and medial pallium of mutants (**Figures 5B, C and S5**), whereas staining was still observed in the marginal zone of the lateral and ventral pallium (**Figures 5B and S5**). Some rare *Reln*^+^ and *Nhlh2^+^* cells were also found ectopically positioned deep within the cortical plate (**Figures 5C**), reminiscent of mice lacking the chemokine *Cxcl12* or its receptors (Borrell and Marín, 2006; Paredes et al., 2006; Trousse et al., 2015). Whole-mount immunostaining followed by tissue clearing and light-sheet imaging confirmed the drastic reduction of Reln^+^ cells in the dorsal cortex, but not the piriform region, of E14.5 mutants compared to wild-type embryos (**Figure 5D**). The absence of CR markers expression in the neocortex of *Gmnc* mutants could either result from mis-specification (CRs would lose the expression of specific genes) or correspond to the absence of CR cells themselves. To formally distinguish between these possibilities and address the origin of remaining Reln^+^ cells in mutants, we performed genetic tracing of hem-derived cells (*Wnt3a^Cre^;Rosa26^tdTomato^*) in control and *Gmnc^-/-^* backgrounds. At E14.5, we observed a drastic reduction in the number of Tomato^+^ cells in the dorsal pallium of *Gmnc* mutants compared to control littermates (**Figure 5E**) or E12.5 mutants (**Figure 4D**), suggesting that the initial incomplete specification of CRs eventually leads to their demise. Of note, CR cells remaining in the lateral and ventral pallium of *Gmnc* mutants did not invade the depleted dorsal pallium at E14.5 (**Figure 5B, E**) or E18.5 (not illustrated), suggesting that self-repulsion (Villar-Cerviño et al., 2013) is not the only mechanism involved in the subtype-specific distribution pattern of CRs. According to *Wnt3a* genetic tracing, a small fraction of these remaining CRs derive from the hem and intermingle with Tomato^-^ CRs most likely derived from the PSB and ThE (**Figures 5E and S3F**).

**Figure 5.**
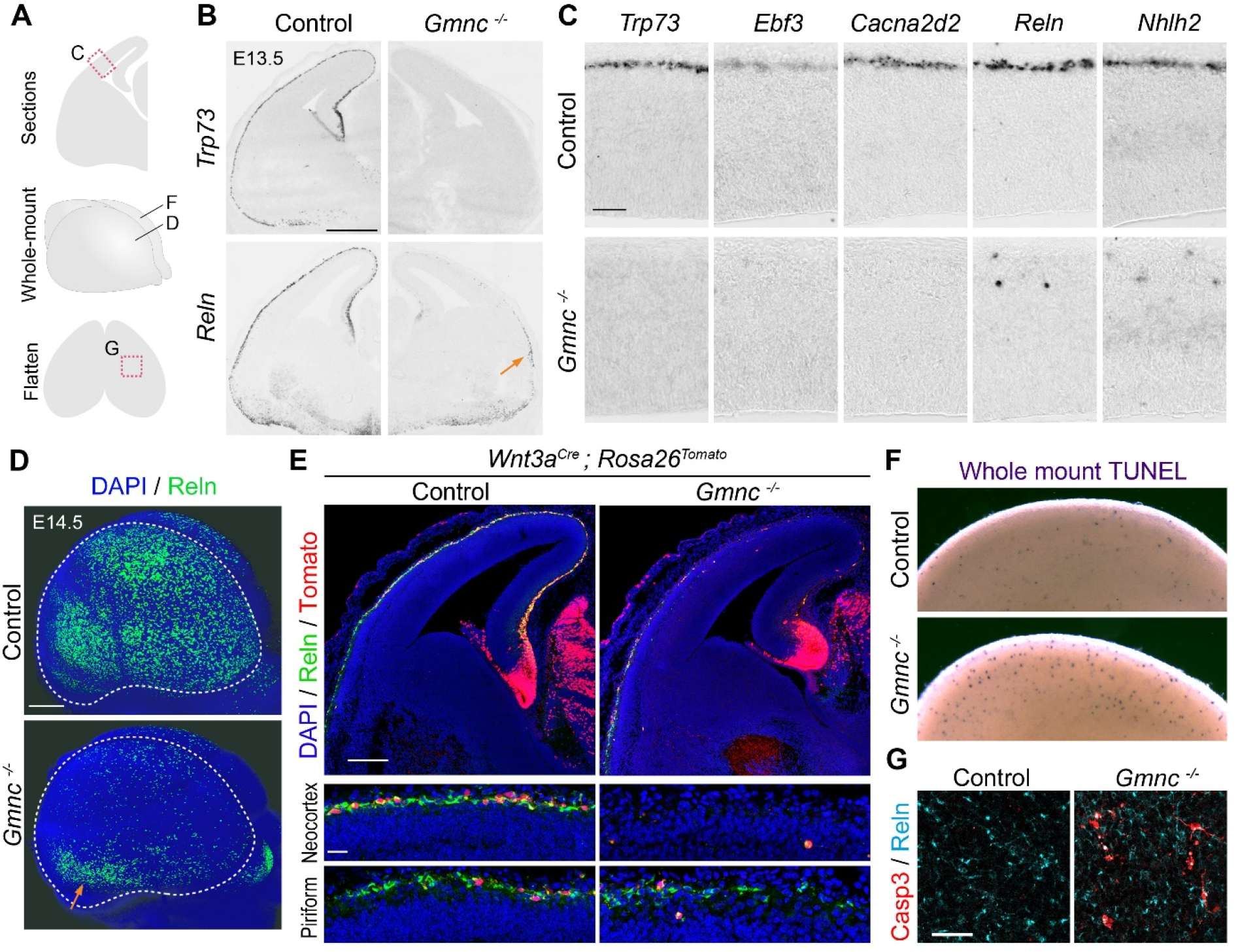
CR cells survival in *Gmnc* mutants. (A) Drawings indicating the regions shown in (B-G). (B) ISH for *Trp73* and *Reln* showing the severe loss of signal in the dorsal cortex of E13.5 *Gmnc* mutants compared to controls. The arrow points to the remaining *Reln^+^* cells in the presumptive piriform cortex. (C) ISH for selected genes specific of either medial (*Trp73, Ebf3* and *Cacna2d2*) or all (*Reln* and *Nhlh2*) CRs subtypes in the dorsal cortex. (D) Light sheet microscopy showing a lateral view of the E14.5 telencephalon following Reln immunostaining (green) in control and *Gmnc* mutants. The arrow points to remaining CRs in the presumptive piriform cortex of mutants. (E) Immunostaining of E14.5 *Wnt3a^Crr^;Rosa26^dTomao^* control or *Gmnc* mutant embryos showing the loss of hem-derived (Tomato^+^, red) and Reln^+^ (green) CRs cells. High magnification (bottom) illustrate the a segment of the neocortex and piriform cortex. (F) Whole-mount TUNEL staining showing increased cell death in the E13.5 dorso-medial cortex of *Gmnc* mutants compared to controls. (E) Surface view of flatten cortex preparations after immunostaining for activated Caspase-3 (red) and Reln (cyan), showing increased CRs apoptosis in E13.5 *Gmnc* mutants relative to wild-type embryos. Scale bars: 500μm in (B and D), 200μm in (E), 50μm in (C and G), 20μm in the high-magnification in (E).

The disappearance of hem-derived CRs in *Gmnc* mutants prompted us to investigate whether cell death occurs following mis-specification. Whole-mount TUNEL staining of E13.5 telencephalic vesicles revealed increased cell death in mutants compared to controls (**Figure 5F**). Consistently, immunostaining for activated-Caspase3 on flat-mount cortical preparations indicated a high number of apoptotic cells, most of which were also Reln^+^, in the marginal zone of mutants compared to controls (**Figure 5G**). We therefore conclude that upon disruption of the multiciliation gene cascade, medial CRs are produced, migrate tangentially, but fail to establish a complete identity and eventually trigger apoptosis.

### Genetic dissection of the multiciliation gene network during CR fate specification

*Gmnc* was previously described as the most upstream element of the multiciliation gene regulatory network, controlling directly or indirectly the expression of all other effectors (Lewis and Stracker, 2021). To test whether such effectors are involved in CR specification, we analysed their contribution using mouse mutants.

*Trp73* is a known player in multiciliation and a direct target of *Gmnc* (Lalioti et al., 2019). In addition, a severe reduction of Reln^+^ cells in the cortical and hippocampal marginal zone of mice lacking *Trp73* has already been reported (Meyer et al., 2004). We subjected E13.5 forebrain sections from *Trp73^-/-^* embryos to ISH using probes directed against pan-CR marker genes *Reln* and *Nhlh2*, as well as medial CR markers *Lhx1* and *Cacna2d2*. We observed a phenotype highly reminiscent of *Gmnc* mutants, consisting in the complete absence of *Lhx1* and *Cacna2d2* expression in the dorsal pallium, the preservation of *Reln*^+^*/Nhlh2*^+^ cells in the marginal zone of the lateral/ventral pallium and the occurrence of ectopic *Reln*^+^ cells deep into the cortical plate (**Figures 6B and S6**). These data suggest that the loss of *Trp73* expression phenocopies that of *Gmnc*, at least for what concerns the establishment and maintenance of hem-derived CR identity. The *Trp73* gene gives rise to several functionally different isoforms (Pozniak et al., 2000; Yang et al., 2000): a transcriptionally active protein, referred to as TAp73 (generated from P1 promoter), and a variant lacking the N-terminal transactivation domain named ΔNp73 (due to usage of an internal promoter in intron 3). Mouse mutants in which a Cre-IRES-EGFP cassette was inserted just after the initiation codon of ΔNp73 display an apparent normal number of GFP^+^ cells at E11.5, that progressively decrease to approximately 50% of controls at P0 (Tissir et al., 2009), suggesting a normal production but abnormal survival of CRs in the absence of ΔNp73. To further study CR fate specification and acquisition of medial identity in these mutants, we performed ISH experiments for pan-CR (*Reln* and *Nhlh2*) and medial CR marker genes (*Lhx1* and *Cacna2d2*). At E13.5, all these genes were specifically expressed in the cortical marginal zone (**Figure 6B**), showing that ΔNp73 is dispensable for the initial acquisition of medial CR fate. Our observations also suggest that hem-derived CRs acquire their medial identity through the *Gmnc*-dependent activation of *Trp73*, and specially point to TAp73 as the responsible isoform.

**Figure 6.**
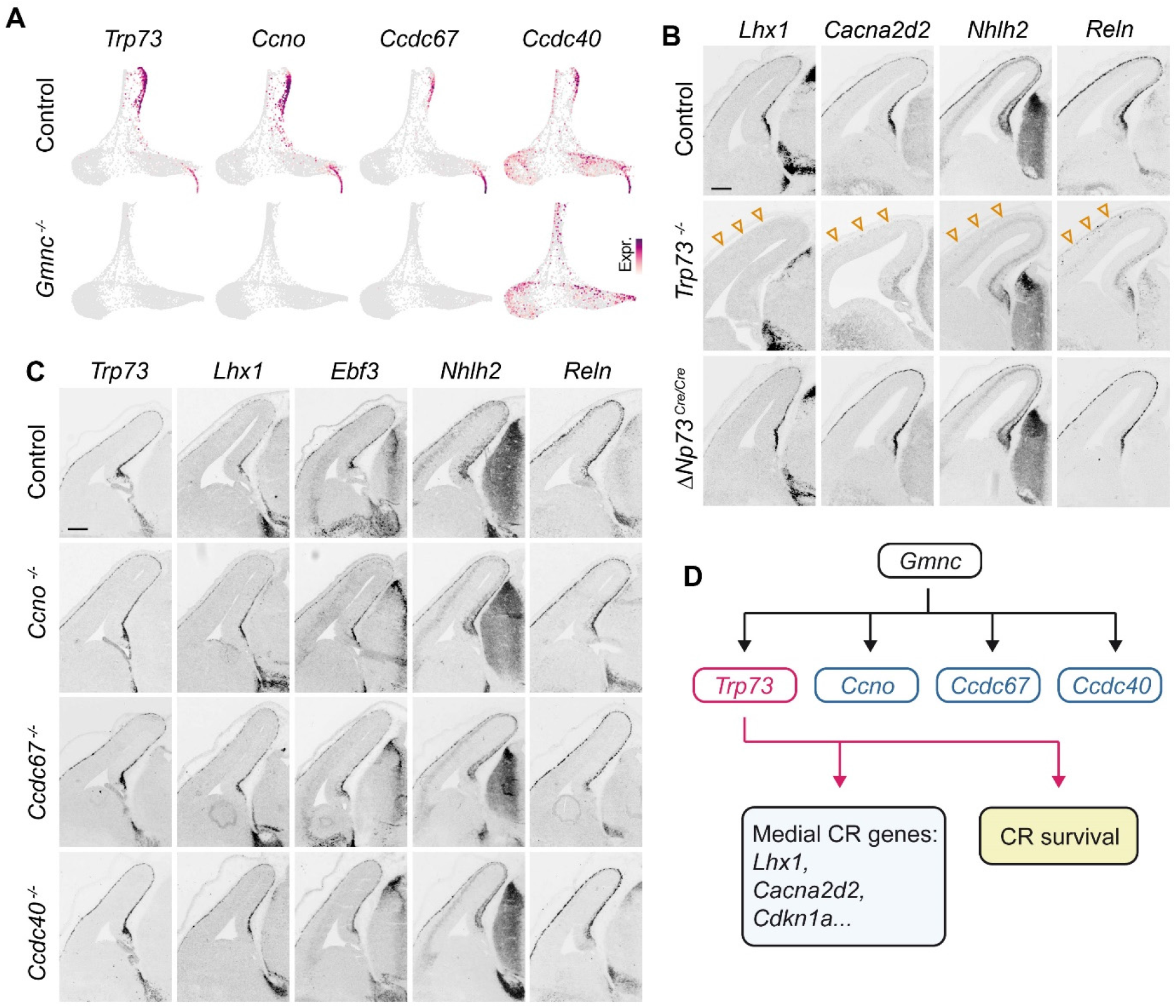
Contribution of multiciliation genes to CR fate specification. (A) SPRING embedding showing the expression of 4 selected genes whose expression is lost, or severely affected, in the CRs lineage of *Gmnc* mutants. (B) ISH for CRs marker genes in E13.5 knock-out embryos with complete deficiency of p73, or with specific inactivation or the ΔNp73 isoform. Arrowheads indicate the absence of staining in the marginal zone of *Trp73* mutants. (C) ISH for CRs marker genes in E13.5 embryos deficient for *Ccno*, *Ccdc67* or *Ccdc40*. Controls and mutants could not be distinguished. (D) Diagram summarizing the proposed contribution of *Gmnc* effectors to CRs differentiation. Scale bars: 200μm.

To further dissect the contribution of *Gmnc* effectors, we thought to determine whether genes previously shown to act downstream *Gmnc* would also be involved in CR fate specification. *Ccno* is a known *Gmnc* target required for multiciliation (Funk et al., 2015; Núnez-Ollé et al., 2017; Terré et al., 2016; Terré et al., 2019). *Ccdc67/Deup1* is also a target of *Gmnc*, essential for the assembly of deuterosome, but its precise function during centriolar amplification and multiciliation remains debated (Mercey et al., 2019; Zhao et al., 2013). Since both *Ccno* and *Ccdc67* were found absent in the CR trajectory of *Gmnc* mutants (**Figure 6A**), we analysed brains obtained from E13.5 embryos lacking either gene to find out that, in both cases, CR markers *Trp73, Lhx1, Ebf3, Nhlh2* and *Reln* were normally expressed in the dorsal pallium (**Figure 6C**).

Finally, we tested whether genes known to control cilia motility and shared by CRs and ChP were required for CR fate. We focussed on *Ccdc40* as we found its expression affected in *Gmnc* mutants (**Figure 6A**). We subjected E13.5 brain sections from *Ccdc40* mutant mice to ISH using probes for *Trp73, Lhx1, Ebf3, Nhlh2 and Reln* to find out that medial CR identity is correctly acquired (**Figure 6C**).

From these experiments, we conclude that *Trp73* is an essential player for CR fate specification downstream *Gmnc*, whereas other tested components of the gene regulatory network are dispensable, at least for the initial acquisition of CR identity (**Figure 6D**). Therefore, despite the significant overlap in their transcriptomic signatures, CRs and ChP differentially interpret the expression of *Gmnc*.

### Expression dynamics and regional cues influence *Gmnc*-induced centriolar amplification

So far, *Gmnc* has only been reported to be a strong inducer of multiciliation. The absence of centriolar amplification in CRs, together with the differential requirement of *Gmnc* downstream effectors in CRs versus multiciliated tissues, raise the question of how the same gene can perform drastically distinct functions in CR and multiciliated lineages.

To determine if the cellular context may influence the ability of *Gmnc* to initiate multiciliation, we implemented *in utero* electroporation experiments. Using this assay, it was shown that the ectopic expression of *Gmnc* in neocortical apical progenitors induces a strong centriolar amplification and retention of electroporated cells at the ventricular surface, reminiscent of ependymal cells, instead of undergoing neuronal differentiation and radial migration towards the pia (Kyrousi et al., 2015; Ortiz-Álvarez et al., 2019; Ortiz Álvarez et al., 2021).

We noticed that the dynamics of *Gmnc* expression differs along ChP and CR differentiation trajectories. In ChP, *Gmnc* expression is initiated in cycling apical progenitor, whereas in CRs it rather starts at the exit of apical progenitor state, and peaks in basal/intermediate progenitors (**Figure 7B**). In order to test whether heterochrony in *Gmnc* expression could restrict its ability to induce multiciliation, we electroporated a LoxP-Stop-LoxP-*Gmnc* construct in *E1-Ngn2^Cre^* embryos to specifically target basal progenitors, and compared to inducing expression in apical progenitors using *Emx1^Cre^* (**Figure 7A**). As expected, electroporations in *Emx1^Cre^* recapitulated that of constitutive *Gmnc* expression in wild-type animals, inducing centriolar amplification and ventricular retention (**Figure 7C**). Most (~60%) electroporated and recombined cells were found in the ventricular zone and displayed large FOP^+^ patches characteristic of centriolar amplification (**Figure 7D**). By contrast, the few cells (~15%) that reached the cortical plate almost systematically showed one small FOP^+^ foci, presumably revealing an absence of centriolar amplification. By contrast, recombined cells in *E1-Ngn2^Cre^* embryos were mostly (~75%) found in the cortical plate and almost never showed centriolar amplification as indicated by FOP immunostaining (**Figure 7E**). We therefore conclude that *Gmnc* expression dynamics, especially the differentiation state at which it peaks, is a key determinant of the cellular response it triggers, consistent with recent findings showing that multiciliation occurs only when *Gmnc* is expressed in cycling progenitors (Ortiz-Álvarez et al., 2019).

**Figure 7.**
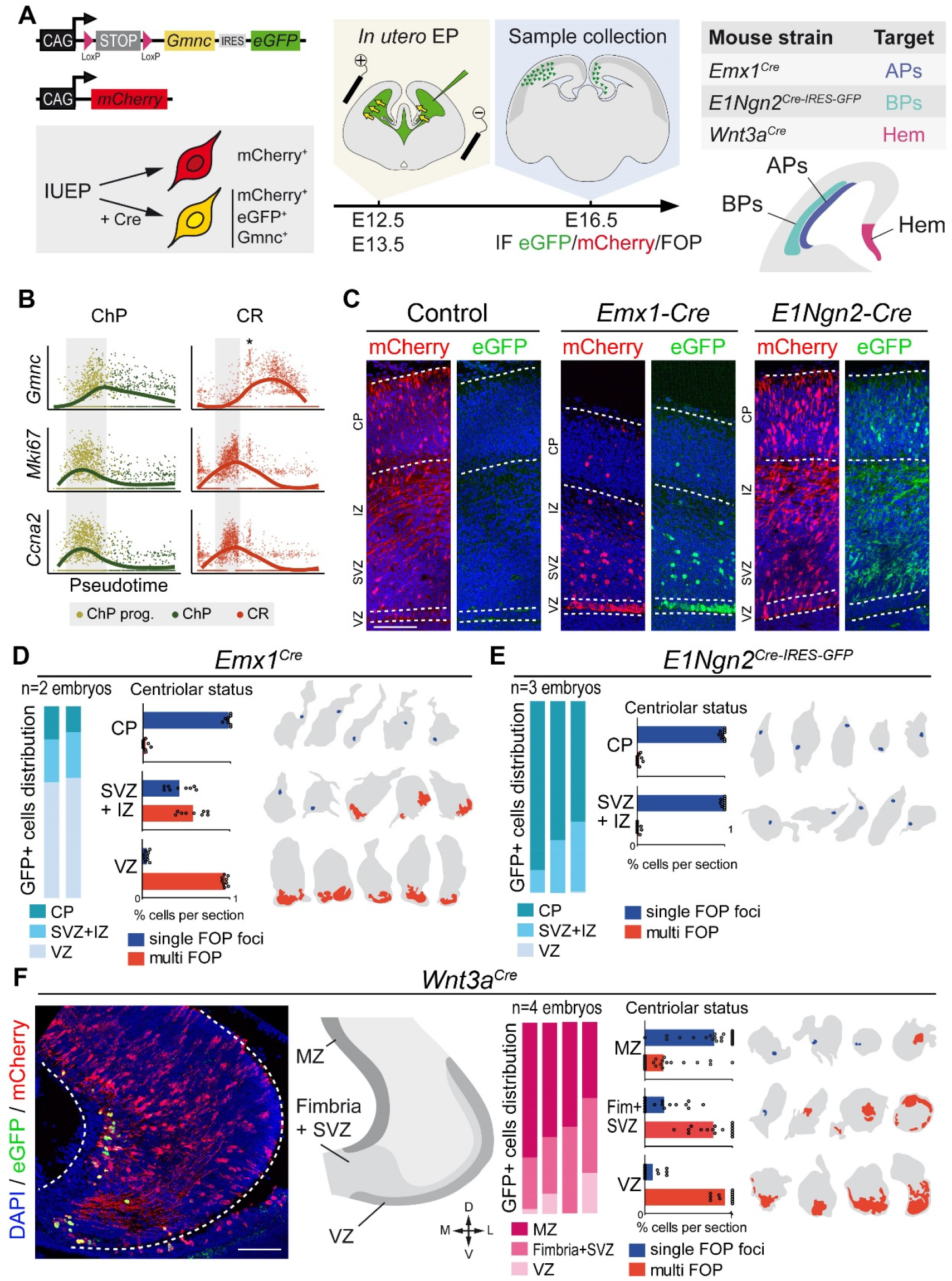
Context-dependent response to *Gmnc* expression. (A) Summary of the experimental design. Plasmids allowing the Cre recombinase-dependent expression of *Gmnc* together with e*GFP*, and the constitutive expression of *mCherry* were combined to monitor recombined *Gmnc*-expressing cells among electroporated cells. *In utero* electroporation experiments were conducted in *Emx1^Cre^, E1Ngn2^Cre^* or *Wnt3a^Cre^* embryos to selectively target apical (APs), basal (BPs) or hem progenitors, respectively. (B) Temporal dynamics of *Gmnc* expression compared to that of *Mki67* and *Ccna2* during ChP (left) or CR (right) differentiation. The area in grey corresponds to S/G2/M phases of the cell cycle, the asterisk indicate cycling BPs. (C) Immunostaining of the dorsal cortex showing the position of electroporated (mCherry^+^, red) and *Gmnc*-expressing cells (eGFP^+^, green). Dashed lines indicate the regions considered for quantifications. (D-E) Quantification of *Gmnc*-expressing cells distribution and centriole amplification after *in utero* electroporation in *Emx1^Cre^* (D) or *E1Ngn2^Cre^* (E) embryos. Cell masks representative of each region are shown next to the histograms; the pattern of FOP staining is represented in blue or red according to the centriolar status. (F) *In utero* electroporation in *Wnt3a^Cre^* embryos. Immunostaining of the hem region showing electoroporated (mCherry^+^, red) and *Gmnc*-expressing cells (eGFP^+^, green). The drawing indicates the regions considered for quantifications. Quantification and representative FOP patterns are shown as in (D-E). 1706 cells were counted from 9 sections of 2 embryos in D, 588 cells from 10 sections of 3 embryos in E, 310 cells from 27 sections of 4 embryos in F. VZ: ventricular zone, SVZ: subventricular zone, IZ: intermediate zone, CP: cortical plate, MZ: marginal zone. Scale bars: 100μm.

Finally, we tested whether spatial cues could also control the response to *Gmnc*. Since apical progenitors of the hem differ from their relatives in the dorsal pallium (see **Figures 1B-D and S1B, C**), we compared the consequence of *Gmnc* expression in these two regions. This was achieved by electroporating the floxed *Gmnc* construct in *Wnt3a^Cre^* embryos (**Figure 7F**). Compared to *Emx1^Cre^* embryos, we found much less ventricular retention of recombined cells, indicating that apical progenitors in the hem respond differently to *Gmnc* expression than their DP counterparts. Similar to both *Emx1^Cre^* and *E1-Ngn2^Cre^* conditions, we found centriolar amplification to occur in the progenitor rather than post-mitotic compartment. Overall, we conclude that both *Gmnc* expression dynamics and hem-specific features contribute to the competence to undergo centriolar amplification or not.

Taken together, our data indicate that components of a well-conserved regulatory gene network involved in centriolar amplification and multiciliation are also essential for the fate specification of CRs derived from medial sources (hem, septum, ThE). We propose that CR fate results from the combination of *Gmnc*-independent processes, sufficient to ensure initial production, tangential migration and *Reln*/*Nhlh2* expression, together with *Gmnc*-dependent mechanisms that enable the acquisition of a complete identity and survival. Our work therefore exemplifies how the co-option of an evolutionary ancient gene module, and redeployment to perform an alternative function, may result in cell type diversification.

## DISCUSSION

In this study we unveil the developmental process involved in the fate specification of medial CRs. Focusing on the major source, the cortical hem, we performed scRNAseq and reconstructed CR differentiation trajectory to unravel a transient expression of the whole gene module controlling multiciliation. Our results, hence uncover the close relationship between medial CR fate specification and acquisition of the multiciliated phenotype of ChP.

The shared expression of *Trp73* by CRs and ChP, and the continuity of their progenitor domains have long suggested a link between their development (Meyer, 2010; Meyer et al., 2019; Roy et al., 2014; Subramanian and Tole, 2009; Tissir et al., 2009). The relationship between CR and ChP fate specification in the telencephalic midline was previously investigated by Imayoshi et al (Imayoshi et al., 2008). In their study, they showed that *Hes1/3/5-mediated* repression of *Neurog2* is required for ChP specification since their combined loss-of-function leads to the overproduction of CRs at the expense of ChP (Imayoshi et al., 2008). Furthermore, ChP to CRs conversion was also observed upon *Neurog2* gain-of-function at early (E9.5) but not late (E11.5) stages of midline patterning, suggesting that progenitors for each lineage are segregated and fully committed by E11.5. Supporting this observation, our data show a clear distinction between CR and ChP progenitors both at E11.5 and E12.5, arguing against the existence of CR/ChP bipotent progenitors at this stage.

The cortical hem is the major source of neocortical CRs in the mouse (Yoshida et al., 2006). It acts as a signalling centre and is considered part of the telencephalon; however, our data clearly highlight the significant transcriptional gap with MP/DP progenitor pools. Interestingly, although the hem, septum and ThE have been studied as independent structures, their molecular identities share some fundamental features which lead to the proposal that they rather represent a continuum referred to as the forebrain-hem system (Roy et al., 2014). Progenitors of the forebrain-hem system express the transcription factors *Id3*, *Otx2*, *Zic2* (Guo and Li, 2019; La Manno et al., 2021; Roy et al., 2014) and their identity is repressed by the pallial transcription factors *Lhx2* and *Foxg1* (Godbole et al., 2017; Godbole et al., 2018; Mangale et al., 2008; Muzio and Mallamaci, 2005; Roy et al., 2014; Vyas et al., 2003). Therefore, the very distinctive transcriptomic signature of CRs, illustrated by their systematic divergence from other cortical excitatory neurons in scRNAseq datasets (Causeret et al., 2021), likely stems from the equally unique identity of their progenitors. In this line, lineage tracing experiments recently demonstrated that neocortical and hippocampal CRs derive from progenitors that are already fate-restricted at E9.5 (Ratz et al., 2022).

Multiciliated ChP and ependymal cells are well conserved among vertebrates (Bill and Korzh, 2014; Fame et al., 2020), and the *Gmnc*-dependent multiciliation regulatory network is also shared by vertebrates (Defosset et al., 2021), suggesting their coevolution. By contrast, CRs have not been described beyond amniotes so far, suggesting they represent a relatively recent evolutionary innovation. We provide strong evidence supporting the existence of CRs in chick, matching previous observations made in lizard and crocodiles (Cabrera-Socorro et al., 2007; Tissir et al., 2003) and confirming that CR identity was already present in the last common ancestor of amniotes at least. Nevertheless, our observation that CR fate specification relies on the multiciliation gene module support a scenario whereby *Gmnc* expression expanded from an ancestral ChP domain into the neighbouring tissue of the forebrain-hem system with a distinct dynamics and was differentially interpreted. Further studies will be necessary to validate such a hypothesis and understand the mechanisms regulating *Gmnc* expression in space and time.

How the same gene module can control two very distinct biological processes remains a puzzling question. During the course of this study, we explored the possibility that the alternative functions of *Gmnc* in CR and multiciliated lineages could be due to a partial co-option of its downstream regulatory network. However, we failed to identify convincing differences in the expression of known *Gmnc* effectors between hem and ChP. Investigating the influence of the cellular context prove more successful, with the demonstration that the cell state and territory in which *Gmnc* is expressed can influence its ability to trigger centriole amplification, which is the first step toward assembling multiple motile cilia (Meunier and Azimzadeh, 2016). The obvious heterochrony of *Gmnc* expression in CRs vs ChP, and its demonstrated impact on centriole amplification corroborate recent work showing that *Gmnc* expression in cortical progenitors favors ependymal fate (Ortiz-Álvarez et al., 2019) and that *Gmnc* induces multiciliation only when expressed in cycling cells (Ortiz Álvarez et al., 2021). By contrast, why hem and DP progenitors are not equally competent remains an open question, although *Foxg1* is an appealing candidate as it was shown to prevent CR fate (Hanashima et al., 2004; Hanashima et al., 2007). We propose that CR specification results from the initial absence of *Foxg1* at the apical progenitor state, followed by the recruitment of the multiciliation regulatory network upon cell cycle exit to progress towards a fully differentiated and viable state without triggering centriole amplification.

The two opposite functional outcomes of the *Gmnc* module in CRs and ChP suggest different target genes are at play in the two lineages. We showed that *Gmnc* mutant phenotype in medial CRs is phenocopied by the loss of *Trp73*, indicating that *Gmnc* function in CR fate specification is mediated by *Trp73*, consistent with previous reports (Amelio et al., 2020; Meyer et al., 2004; Meyer et al., 2019; Yang et al., 2000). Interestingly, *Trp73* is a key player in ependymal specification, maturation and organization (Fuertes-Alvarez et al., 2018; Gonzalez-Cano et al., 2016; Medina-Bolívar et al., 2014). It is also a well-known regulator of multiciliation through its direct control on the transcription of key effector genes (Marshall et al., 2016; Nemajerova et al., 2016). Although further work will be required to identify *Trp73* targets in CRs vs ChP, it is tempting to speculate that the co-option of *Trp73* by CR cells has been followed by the acquisition of new target genes, leading to a functional divergence from ChP.

CRs have a transient lifetime, previously prompting us to postulate that they could be fated to die (Causeret et al., 2018). In this frame, our observation that misspecified CRs undergo premature apoptosis in *Gmnc* mutants suggests that acquisition of their complete identity is required to maintain them alive. Such hypothesis is supported by the demonstrated role of ΔNp73 in the survival of CRs (Tissir et al., 2009). It also raises the question of PSB-derived CRs which never express *Gmnc* or *Trp73*. One could speculate that their similarity with mis-specified medial CRs in *Gmnc* mutants foretells a short lifetime, supporting further investigations to precisely determine the survival curve of PSB-derived CRs.

CRs represent a rare but very important cell type in the developing neocortex, involved in multiple aspects of its morphogenesis, and proposed to contribute to its tremendous expansion and complexification in mammals (Causeret et al., 2021; Goffinet, 2017). We believe that our work perfectly exemplifies how the co-option of a single gene, followed by the refunctionalisation of its downstream regulatory network, may result in the invention of a novel cell type, and how cell type diversification can occur by dramatic leap rather than progressive divergence.

## MATERIALS AND METHODS

### Animals

The following mouse lines were used and maintained on a C57BL/6J background: *Gmnc*^-/-^ (*Gmnc^tm1.1Strc^*) (Terré et al., 2016), *Trp73*^-/-^ (*Trp73^tm1Fmc^*) (Yang et al., 2000), *Ccdc67*^-/-^ (*Deup1^tm1.1(KOMP)Vlcg^*) (Mercey et al., 2019), *Ccno*^-/-^ (*Ccno^tm1.1(KOMP)Vlcg^*) (Núnez-Ollé et al., 2017), *Ccdc40*^-/-^ (*Ccdc40^lnks^*) (Becker-Heck et al., 2011), *Foxj1^CreERT2^* (*Foxj1^tm1.1(cre/ERT2/GFP)Htg^*) (Muthusamy et al., 2014), *PGK^Cre^* (*Tg(Pgk1-cre)1Lni*) (Lallemand et al., 1998), *Emx1^Cre^* (*Emx1^tm1(cre)Krj^*) (Gorski et al., 2002), *ΔNp73^Cre^* (*Trp73^tm1(cre)Agof^*) (Tissir et al., 2009), *Wnt3a^Cre^* (*Wnt3a^tm1(cre)Eag^*) (Yoshida et al., 2006), *E1-Ngn2^Cre^* (*Tg(Neurog2-cre/GFP)1Stoy*) (Berger et al., 2004), *Rosa26^YFP^* (*Gt(ROSA)26Sor^tm1(EYFP)Cos^*) (Srinivas et al., 2001) and *Rosa26^tdTomato^* (*Gt(ROSA)26Sor^tm9(CAG-tdTomato)Hze^*) (Madisen et al., 2010). All animals were handled in strict accordance with good animal practice, as defined by the national animal welfare bodies, and all mouse work was approved by either the French Ministry of Higher Education, Research and Innovation as well as the Animal Experimentation Ethical Committee of Université Paris Cité (CEEA-34, licence numbers: 2020012318201928 and 2018020717269338), or the institutional ethics committee from Universidad de Leon and Junta de Castilla y Leon (approval OEBAULE-018-2017 and OEBA-ULE-018-2021).

### scRNAseq

Four *PGK^Cre^*; *Rosa26^YFP^* E11.5 embryos and four *Wnt3a^Cre^*; *Rosa26^tdTomato^* E12.5 embryos were collected in HBSS on ice to generate the wild-type reference dataset. *Gmnc*^-/-^ cells were obtained from four pooled E12.5 mutant embryos. For each dataset, a region spanning the dorsal pallial midline (see Figure 1A) was dissected on both hemispheres and dissociated using the Neural Tissue Dissociation Kit (P) (Miltenyi Biotec) on the gentleMACS Octo Dissociator following the manufacturer’s instructions. Remaining clumps of cells and debris were removed via two rounds of centrifugation for 3 min at 200 g followed by filtration through 30 μm cell strainers (Miltenyi Biotec). To ensure all remaining cells aggregates were dissociated, we finally pipetted the suspension through Gel Saver tips (QSP) and proceeded to a last filtration through a 10 μm cell strainer (pluriSelect). Cell viability and concentration were assessed using a MACSQuant flow cytometer (Miltenyi Biotec). Approximatively 20000 cells were used as input on the 10X Genomics Chromium Controller for a targeted cell recovery of 10000 cells per library. The entire procedure, from the sacrifice of the pregnant female until loading the controller, was achieved in 2h. For the wild-type dataset, two Single Cell 3’ Library and Gel Bead Kit v3 libraries were produced and sequenced on a NovaSeq 6000 sequencer at a total depth of 800 million reads. For the mutant dataset, one library was produced and sequenced at a depth of 750 million reads. Raw sequencing reads were processed to counts matrix with Cell Ranger version 3.1.0 using default parameters and the mm10 mouse genome reference to which the sequenced of *YFP* and *tdTomato* were added. All statistical analyses were performed under R version 4.1.2.

### Cell filtering

Each library was processed independently to extract high-quality cells from the raw count matrix. Cells were filtered based on the percentage of UMI associated with mitochondrial transcripts and number of detected genes, to keep only those within a three median absolute deviation (MAD) around the population median for these metrics. We further excluded cells having a UMI number above 3 MADs of the population median. Potential doublets were removed using Scrublet (Wolock et al., 2019). Dimensionality reduction was performed using the SPRING tool (Weinreb et al., 2018). To combine wildtype and mutant datasets into the same reduced space, we projected the mutant cells onto control cells using the dedicated algorithm implemented in SPRING. Counts values were library size normalised and scaled using Seurat (v 4.0.5) (Hao et al., 2021).

### Broad clustering and cell type annotation

For each dataset, we first apply coarse Louvain clustering and use known marker gene to identify the major classes of radial glia progenitor, and neuronal cells. Wild-type neuronal cells were extracted and separated into two lineages during a second round of Louvain clustering. Belonging to CRs or pallial neurons lineages was readily identified based on the exclusive expression of the transcription factors *Trp73* or *Foxg1*.

To further resolve the diversity among wild-type progenitors and ChP populations, we first performed topic modelling using fastTopic (v.5.59) (Carbonetto et al., 2021; Dey et al., 2017). Then, progenitors were grouped into 6 clusters by applying partition around medoids to the loading of the 6 topics which best captured progenitor domain identities. Finally, top marker genes for each inferred cluster were used to validate their annotation by mapping their position on tissue using ISH.

In order to annotate the mutant dataset, we first extracted progenitor cells and transferred the wild-type ISH-validated cluster annotation using SeuratV4 functions “FindTransferAnchors” and “TransferData”. We then extracted neuronal cells and identified the most mature cells after fitting a principal curve to their SPRING coordinates. These cells were then subject to iterative clustering using scrattch.hicat (v0.0.16) (Tasic et al., 2016) to identify the 3 population corresponding respectively to the hem- and ThE-derived CRs and pallial neurons. Finally, annotation of the intermediate states cells was predicted using FateID (v0.1.9) (Herman et al., 2018) seeded with the 3 clusters of mature neurones found in this dataset.

### Trajectories

For each neuronal differentiation trajectory, a pseudotime axis was fitted in two steps. First, we extracted apical progenitors belonging to the cortical hem or medial pallium cluster. These progenitors were then aligned along their cell cycle axis, from G1 to M phase using the Revelio package (v0.1.0). In parallel, a principal curve was fitted on the SPRING coordinates of the two neuronal lineages cells to capture the panneuronal differentiation axis. In a second step, the two ordering of progenitors and neurons were concatenated into the same pseudotime axis, with progenitors arbitrarily allocated to the first 1/3 of the full trajectory. With this approach, differentiation trajectories therefore start at the G1 phase in apical progenitors.

Differentially expressed genes along pseudotime between the two trajectories were identified using the likelihood ratio test implemented in the function ‘differentialGeneTest’ of the package Monocle (v2.22.0). Trajectory-specific enrichment was determined by computing the area between the two curves as performed by Monocle. Genes with similar smoothed trends were grouped into 6 clusters using partition around medoids. Finally, gene list functional enrichment analysis was performed using gprofiler2 (v2_0.2.1) (Kolberg et al., 2020).

### Cloning

*Gmnc* coding sequence (CDS) was inserted into a pCAGGs-Lox-STOP-Lox-IRES-GFP (pCAG-LSL) as follows. First, *Gmnc* CDS was amplified by PCRs from a pCAGGS-Gmnc plasmid (Kyrousi et al., 2015) using Phusion DNA polymerase (Thermo Scientific F530S) and primers carrying XhoI or SalI restriction sites. Then, DNA digestion of the pCAG-LSL and *Gmnc* cDNA was performed with XhoI (Thermo Scientific FD0694) and SalI (Thermo Scientific FD0644) and the *Gmnc* fragment inserted in the backbone by enzymatic ligation using T4 ligase (Thermo Scientific EL0014). Plasmid amplification was performed in One Shot™ TOP10 competent E. coli (Invitrogen C404010) and purification was achieved using an Nucleobond Xtra Maxi EF kit (Macherey-Nagel 740424.10). Plasmids were verified by restriction and sequencing to confirm the correct insertion.

### *In utero* electroporation

E12.5 or E13.5 pregnant females (wild-type or transgenic) were anaesthetised with Isoflurane (4% induction, 2% during the surgery) and subjected to abdominal incision to expose the uterine horns. A combination of pCAG-LSL-Gmnc-GFP and pCAGGs-mCherry (Arai et al., 2019) at a final concentration of 1 μg/μl and mixed with Fast Green was injected into the lateral ventricles of embryos with a glass capillary. A train of five pulses (30 V, 50 ms, 1Hz) was applied to injected embryos through the uterus using a NEPA21 electroporator (Nepagene). Uterine horns were repositioned into the abdominal cavity, and the abdominal wall and skin were sutured. Embryos were harvested 3days later.

### Tissue processing

Embryos were staged, starting from midday of the vaginal plug considered as embryonic day 0.5 (E0.5). After collection and dissection in ice-cold PBS, embryos were immediately fixed by immersion in 4% paraformaldehyde, 0.12 M phosphate buffer (PB) (pH 7.4), at 4°C for 2 h for E11.5-E12.5 to 3 h for E13.5. Cryoprotection was achieved by overnight incubation in 10% sucrose, PB at 4°C, before embedding in 7.5% gelatine, 10% sucrose in PB, and freezing by immersion in isopentane cooled at −55°C. 20 μm coronal sections were obtained with a Leica CM3050 cryostat and collected on Superfrost Plus slides (Menzell-Glasser).

### Tissue clearing

We performed whole head clearing according to the advanced CUBIC protocol (Susaki et al, 2015). Tissues were fixed with 4% paraformaldehyde in PBS, for 2h30 at 4°C, and washed 3 times in PBS 1X for 1 hours. For the delipidation step, tissues were incubated in the Sca*l*eCUBIC-1 reagent at RT for 1 week followed by 6 washes in PBS 1X for 1h. After permeabilisation in PBS 0.2% Gelatin, 0.5% Triton-X100, tissues were incubated with primary antibodies for 1 week. Excess of antibodies were removed during 6 washes in PBS 1X for 1h. Secondary antibodies were incubated with TO-PRO-3 for 3 days. All steps were performed at RT with agitation. Brains were finally cleared in the aqueous solution of Sca*l*eCUBIC-2 reagent.

### Immunostaining

The following primary antibodies were used: chick anti-GFP (Aves Labs GFP-1020, 1:1000), rabibit anti-Tomato (Clontech 632496 1:500), mouse anti-Ebf3 (Abnova H00253738-M05 1:1000), mouse anti-Reln (Millipore MAB5364 1:300), rabbit anti-Trp73 (Cell signaling 14620 1:250), rabbit anti-Tbr2 (Abcam ab216870, 1:300), rabbit anti-Foxg1 (Abcam ab18259 1:1000), rabbit anti-Caspase 3 (Cell Signaling 9664, 1:400), mouse anti-FOP (Abnova H00011116-M01 1:500), anti-Arl13b (NeuroMab 75-287, 1:500), mouse anti-p21 (1:100, Santa Cruz Biotechnology, sc-6246). The following secondary antibodies were obtained from Jackson ImmunoResearch: donkey anti-chick Alexa-488 (1:1000), donkey anti-mouse Cy5 (1:500), donkey anti-rabbit Cy3 (1:700) and donkey anti-rabbit Cy5 (1:500). DAPI (1 μg/ml) was used for nuclear staining. Slides were mounted in Vectashield (Vector).

### *In situ* hybridisation

For each gene of interest, a DNA fragment (typically 500-800 bp) was amplified by PCRs from an embryonic brain cDNA library using Phusion polymerase (Thermo). Primers for mouse *Cacna2d2*, *Chl1, Ebf3, Gm27199, Grm1, Lhx1* and *Unc5c* are described elsewhere (Moreau et al., 2021) Table S1. The promoter sequence of the T7 RNA polymerase (GGTAATACGACTCACTATAGGG) was added in 5’ of the reverse primer. Alternatively, for mouse *Foxg1*, *Reln*, *Trp73* and *Wnt3a*, a plasmid containing part of the cDNA was linearised by enzymatic restriction. Antisense digoxigenin-labelled RNA probes were then obtained by *in vitro* transcription using T7, T3 or SP6 RNA polymerase (New England Biolabs) and digRNA labelling mix (Roche). *In situ* hybridisation was carried out as previously described (Schaeren-Wiemers and Gerfin-Moser, 1993) using a hybridisation buffer composed of 50% formamide, 5×SSC, 1×Denhardt’s, 10% dextran sulfate, 0.5 mg/ml yeast RNA and 0.25 mg/ml herring sperm DNA. Probes were detected using an anti-digoxigenin antibody coupled to alkaline phosphatase (Roche) and NBT/BCIP (Roche) as substrates. Slides were mounted in Mowiol.

### Whole mount TUNEL staining

Whole mount TUNEL was performed as described previously (Causeret et al., 2011) using the Apoptag Peroxidase in situ Apoptosis Detection kit (Millipore) according to the manufacturer’s instructions. Embryos were digested with 10 μg/mL proteinase K for 3 min, followed by 1h incubation with the terminal deoxynucleotidyl transferase at 37°C. Cell death was revealed using the alkaline phosphatase-conjugated anti-digoxigenin antibody (Roche) and NBT/BCIP (Roche) as a substrate.

### Image acquisition

*In situ* hybridisation images were obtained using a Hamamatsu Nanozoomer 2.0 slide scanner with a 20× objective. Immunofluorescence images were acquired using a Leica SP8 confocal microscope with a 40× objective. 3D image acquisition of optically cleared samples was performed using a Zeiss Lightsheet Z1 microscope with a 20X objective. Images were analysed using ImageJ software and figures composed with Adobe Photoshop.

### Resources availability

Raw scRNAseq data, processed data and metadata will be deposited on GEO upon publication. Comprehensive R codes can be found at https://github.com/MatthieuXMoreau/hemCR. Further information and requests for resources and reagents should be directed to and will be fulfilled by the lead contact, F. Causeret.

## ACKNOWLEDGEMENTS

The authors wish to thank Marine Luka and the LabTech Single-Cell@Imagine for their support in implementing scRNAseq experiments, as well as Cecile Masson and all the staff from the Imagine genomic and bioinformatics core facilities for advice and technical assistance. We are grateful to Francesco Carbone and Jérôme Flatot for implementing the Shiny App server, and Institut Français de Bioinformatique for providing access to the R-Studio service of the IFB-core cluster. We acknowledge the histology and cell imaging facilities of the Structure Fédérative de Recherche Necker (Inserm US24, CNRS UAR3633) and the NeurImag Imaging core facility of IPNP. We thank Leducq establishment for funding the Leica SP8 confocal/STED 3DX system at IPNP and Association pour la Recherche sur le Cancer for funding the Nanozoomer slide scanner at SFR Necker. We are grateful to Sébastien Dupichaud for his help with light sheet microscopy. We acknowledge the Imagine Institute LEAT and IPNP animal facility for animal care, especially the zootechnicians in charge of our colonies. We thank Sigolène Meilhac and Anna-Katerina Hadjantonakis for providing *Ccdc40/Inks* mutants. We are grateful to all members of the Pierani and Spassky labs for stimulating and helpful discussions, and Pierre Billuart for critical reading of the manuscript.

## AUTHOR CONTRIBUTIONS

Conceptualization: MXM, NS, AP, FC

Methodology: MXM, YS, FC, NS, AP, FC

Software: MXM, FC

Validation: MXM, YS, VE, BB, ED, TDG, NS, FC

Formal analysis: MXM

Investigation: MXM, YS, VE, BB, ED, TDG, NS, FC

Resources: AO, LMA, MMM, MCM, NS

Data curation: MXM, FC

Writing - original draft: MXM, FC

Writing - review & editing: MXM, NS, AP, FC

Visualization: MXM, YS, VE, BB, ED, NS, FC

Supervision: FC

Project administration: FC

Funding acquisition: AP, FC

## SUPPLEMENTARY MATERIAL

**Figure S1.**
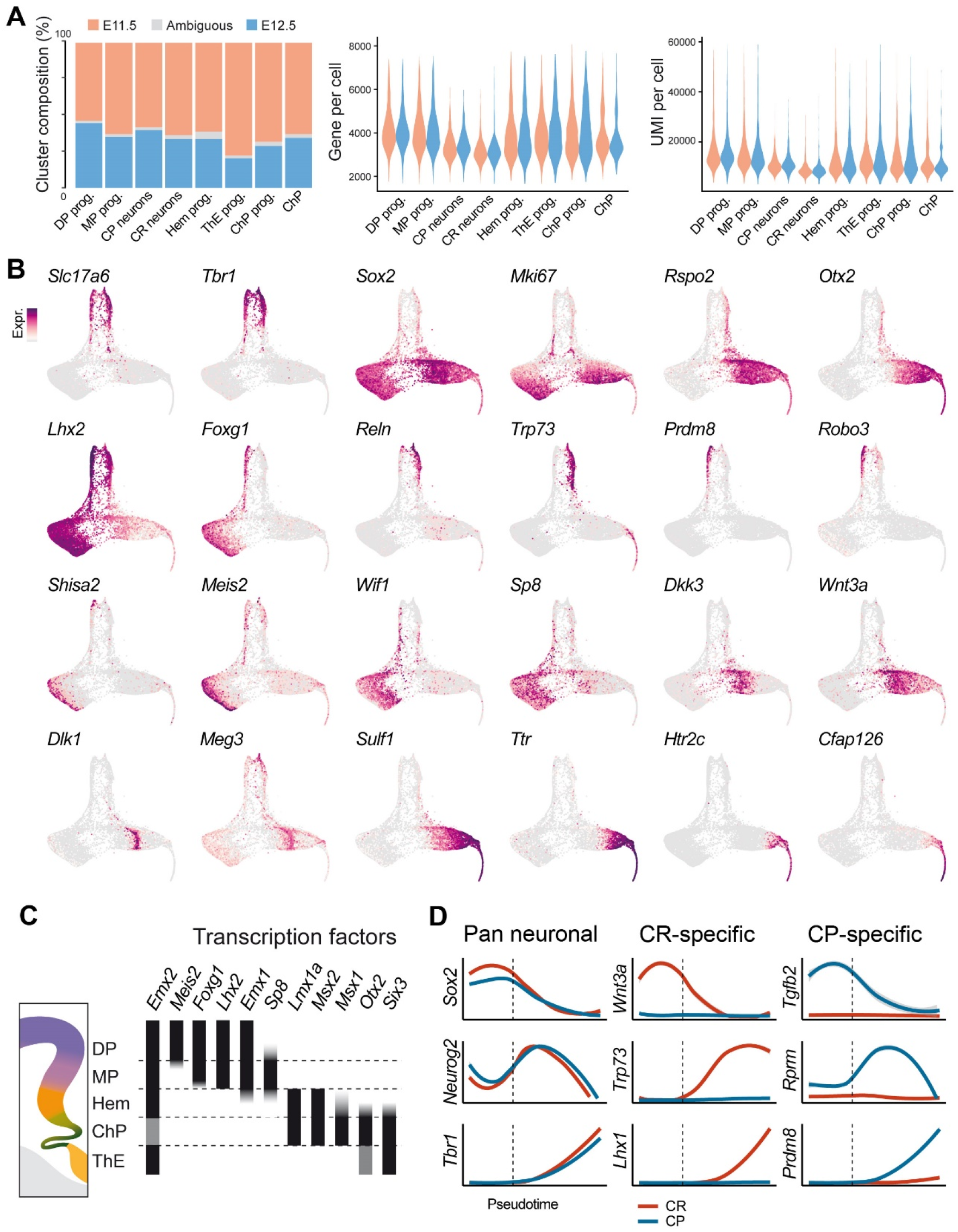
Additional gene expression patterns. (A) Estimated contribution of cells collected at E11.5 or E12.5 to each cluster, and comparison of the number of genes and UMI per cell at each age. (B) Expression of selected genes on the SPRING dimensionality reduction used to annotate the dataset (C) Illustration of the expression of selected transcription factors in progenitor domains (D) Smoothed expression dynamics of selected genes along CRs (red) and CP neurons (blue) differentiation. The dashed line represents the exit of apical progenitor state.

**Figure S2.**
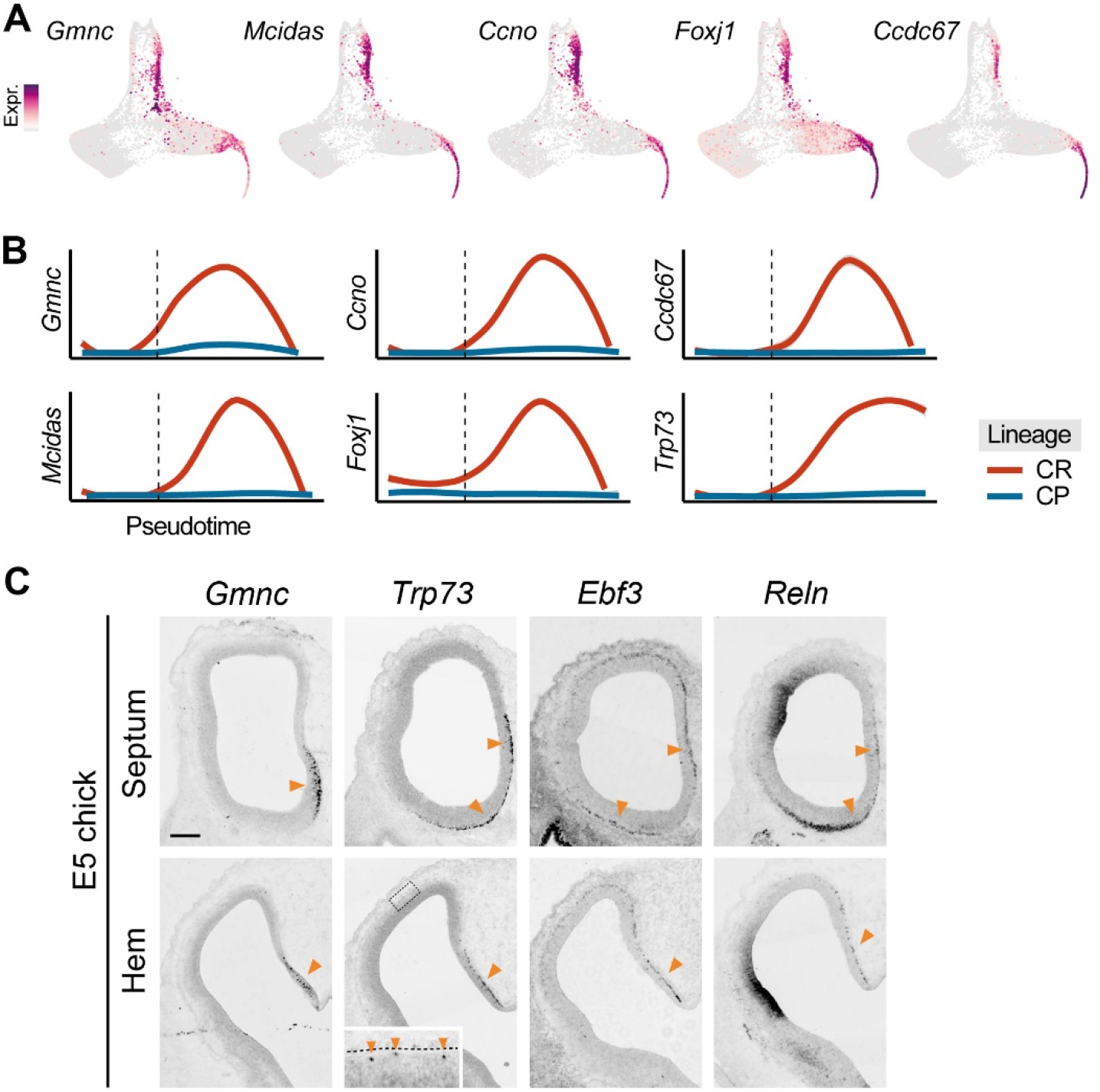
Expression of multiciliation genes. (A) Expression of selected multiciliation genes on the SPRING dimensionality reduction used to annotate the dataset (B) Smoothed expression dynamics of multiciliation genes along CRs (red) and CP neurons (blue) differentiation. The dashed line represents the exit of apical progenitor state. (C) ISH for *Gmnc, Trp73, Ebf3* and *Reln* on coronal sections of E5 chick embryos at the level of the septum and hem. Arrowheads point to signal in the marginal zone. The high magnification for *Trp73* illustrate the few positive cells found in the dorsal cortex, far from the hem. Scale bar: 200μm

**Figure S3.**
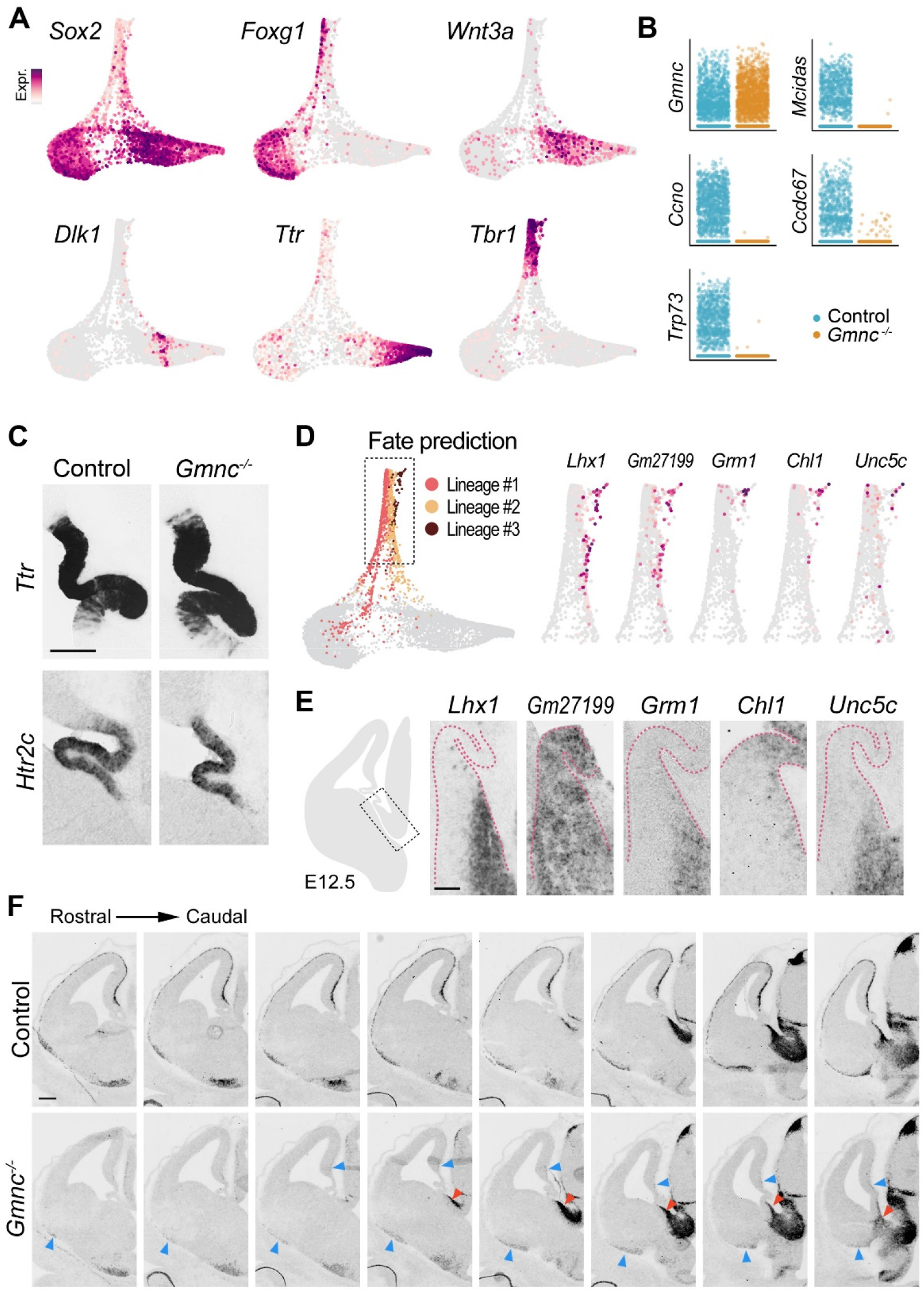
Additional characterization of *Gmnc* mutants. (A) Expression of selected genes on the SPRING dimensionality reduction of the *Gmnc* mutant dataset. (B) Comparative expression of multiciliation genes in control (blue) and *Gmnc* mutant (orange) cells. Mutant *Gmnc* transcripts lack exons 3 and 4 but retain the 3’UTR and are therefore detected by scRNAseq. (C) ISH for ChP-specific genes on coronal sections of E12.5 control or mutant embryos. (D) Neuronal lineages identified in the *Gmnc* mutant dataset. High magnifications (right panel) illustrate the expression of selected markers of the small population that was excluded from subsequent analyses. (E) ISH for genes shown in (D) on coronal sections of the E12.5 ThE from wild-type embryos. (F) ISH for *Lhx1* on coronal sections of the E12.5 forebrain from wild-type and *Gmnc* mutant embryos. Red arrowheads point to the ThE, blue arrowheads indicate the rare *Lhx1^+^* cells observed in the telencephalic marginal zone of *Gmnc* mutants. Scale bars: 50μm in (E), 200μm in (F).

**Figure S4.**
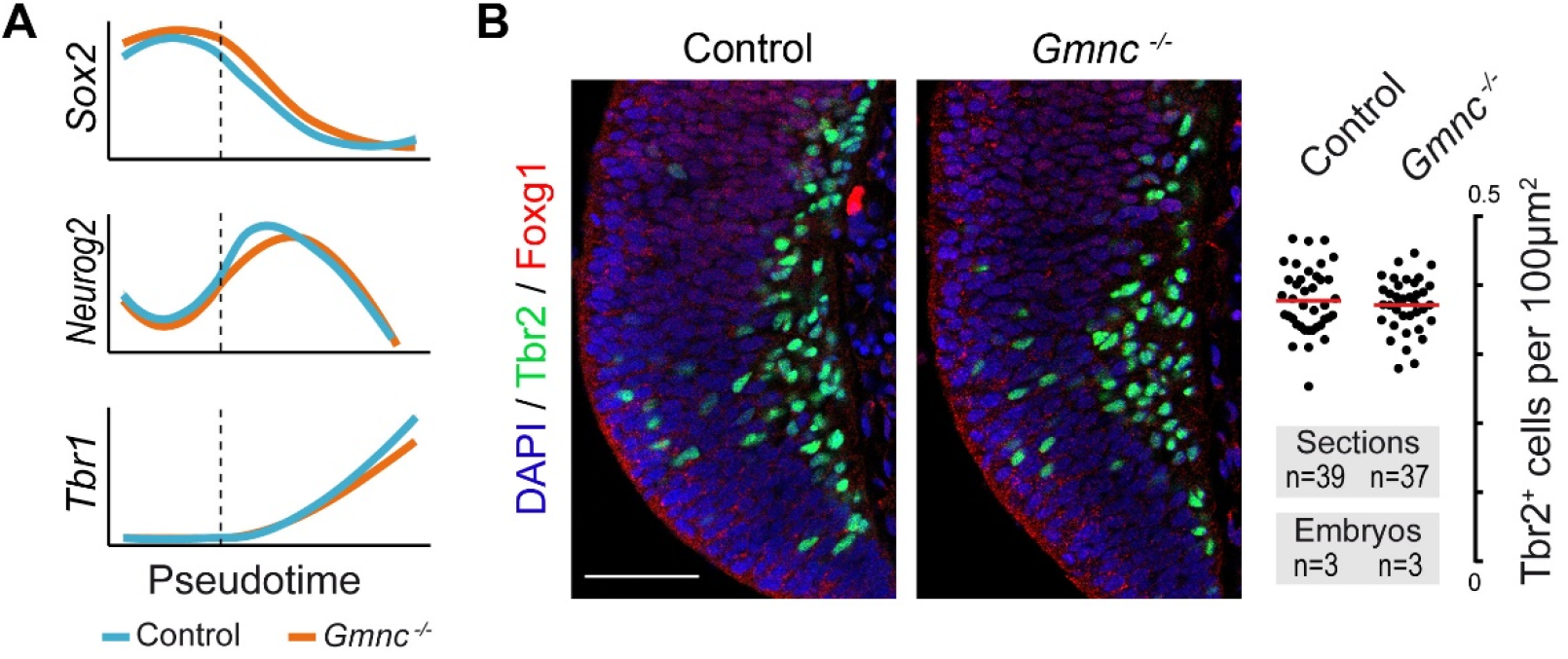
Neurogenesis in *Gmnc* mutants. (A) Temporal dynamics of genes expressed in apical progenitor (*Sox2*), basal progenitors (*Neurog2*) or differentiated neurons (*Tbr1*) during CRs differentiation in control (blue) and *Gmnc* mutant (orange) datasets. The dashed lines correspond to the exit of apical progenitor state. (B) Immunostaining on coronal sections of E12.5 control or *Gmnc* mutant embryos showing basal progenitors and early neurons (Tbr2^+^, green) in the hem region identified by the absence of Foxg1 (red) staining. Quantification of the density of Tbr2^+^ cells revealed no significant differences between control and mutants. Scale bar: 50μm.

**Figure S5.**
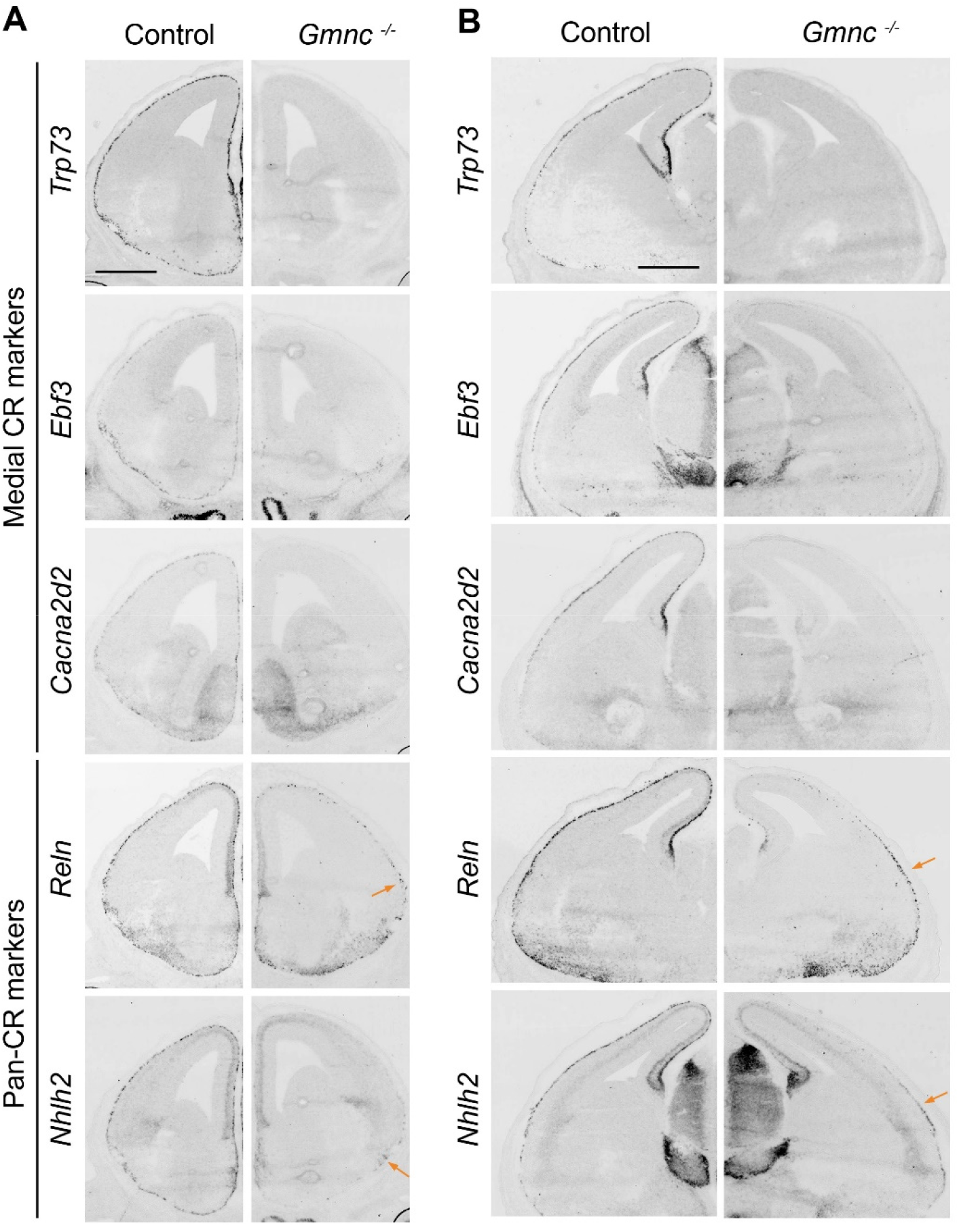
Expression of CR marker genes in *Gmnc* mutants. (A) ISH for marker genes of medial CRs (*Trp73, Ebf3, Cacna2d2*) or all CRs subtypes (*Reln, Nhlh2*) on coronal section at rostral levels of E13.5 *Gmnc* mutants and controls. (B) same as A at caudal levels. Arrows point to remaining CRs in lateral and ventral regions of mutants. Scale bar: 500μm.

**Figure S6.**
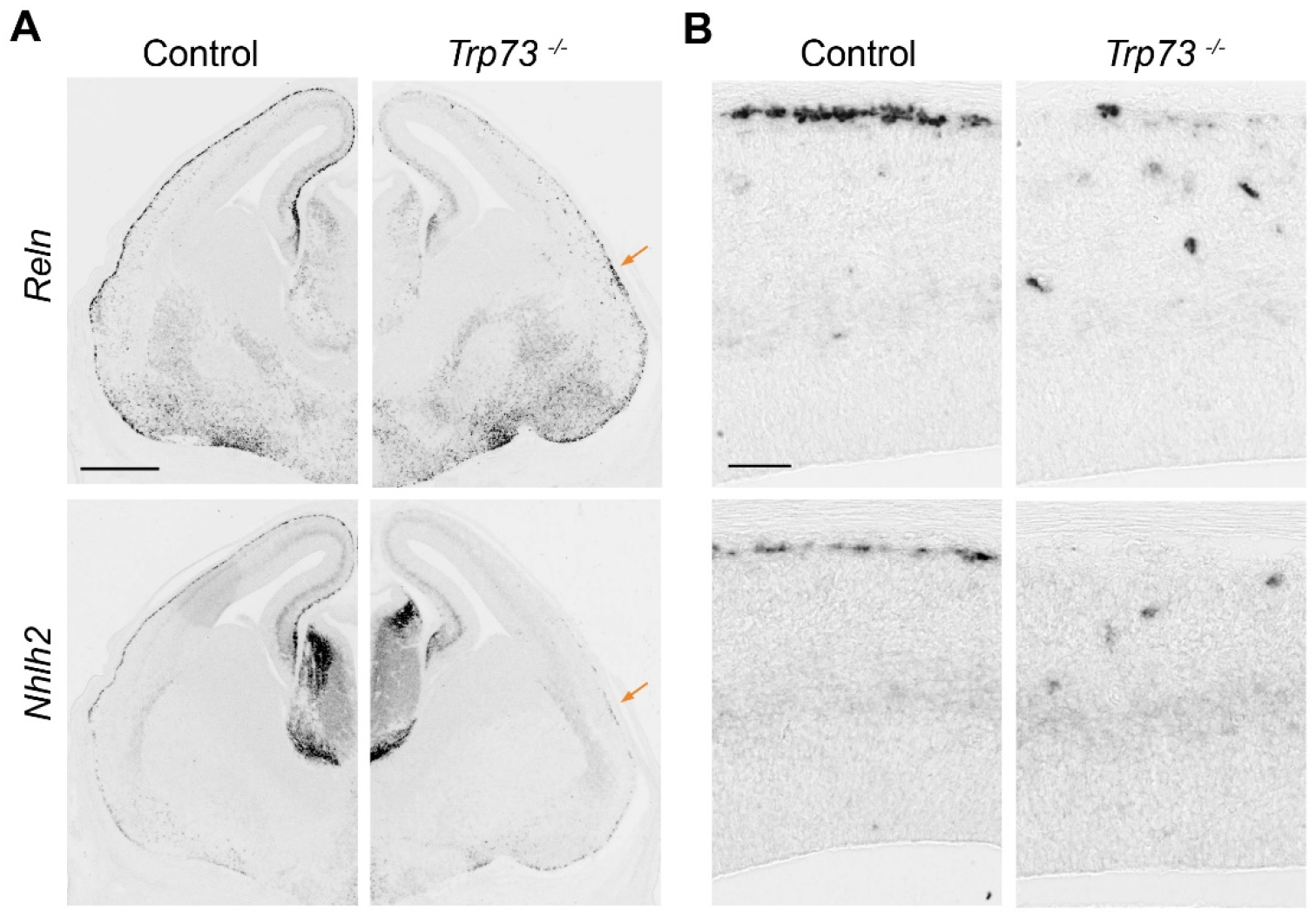
Expression of CR marker genes in *Trp73* mutants. (A) ISH for *Reln* and *Nhlh2* on coronal section of E13.5 *Trp73* mutants and controls. The arrow points to the few remaining CRs in lateral and ventral regions. (B) High magnification of the dorsal cortex showing the few ectopic CRs found deep into the cortical plate in mutants instead of their normal superficial position. Scale bars: 500μm in A, 50μm in B.

**Table S1.**
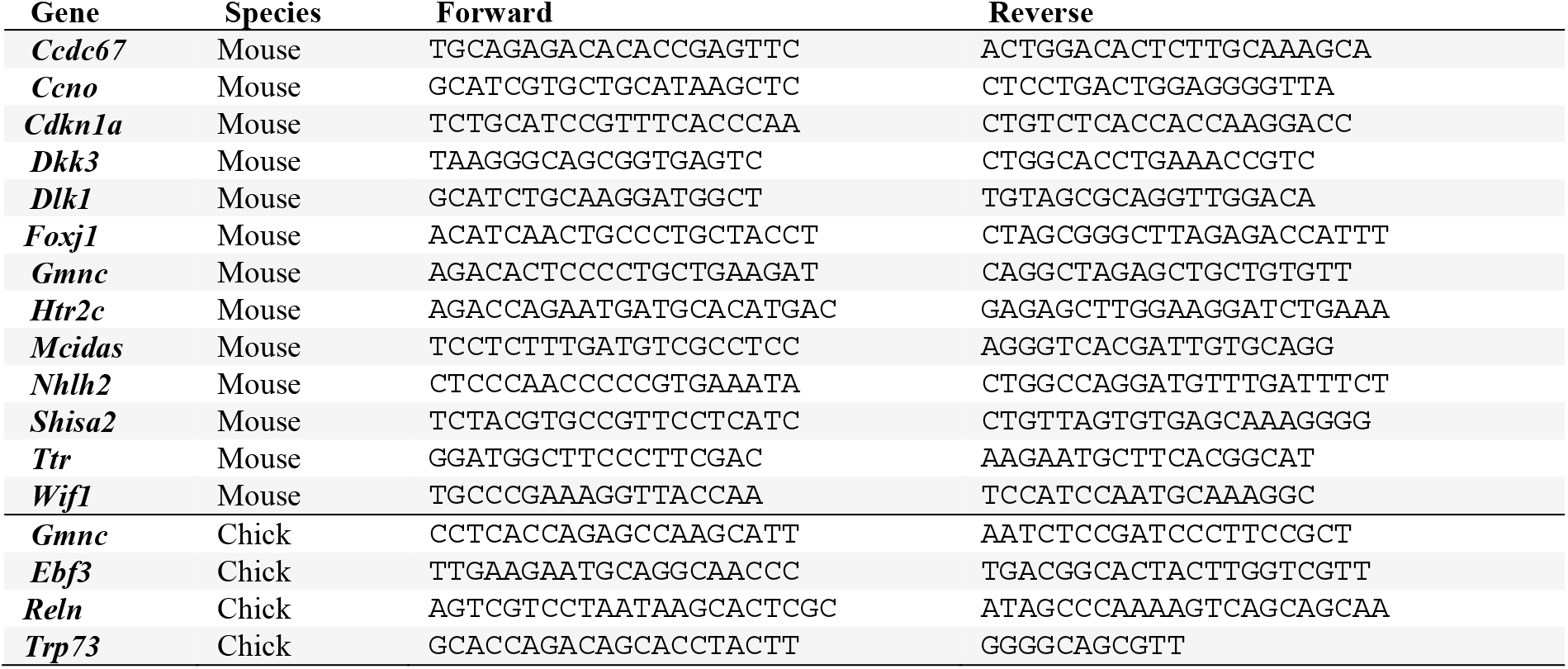
Primers for ISH probes.

